# Caribbean octocoral communities: finding the forest for the trees?

**DOI:** 10.1101/2025.01.05.631368

**Authors:** Howard R Lasker, Lorenzo Bramanti, Peter J. Edmunds, John F. Girard, Nolwenn Pages, Kaitlyn Tonra, Christopher D. Wells, Katell Guizien

**Affiliations:** University at Buffalo, Buffalo, New York, United States of America; Sorbonne Université, CNRS, LECOB-Observatoire Océanologique de Banyuls, France; California State University, Northridge, United States of America; University of Rhode Island, Kingston, Rhode Island, United States of America; Oregon State University, Corvallis, Oregon,United States of America; Bowdoin College, Brunswick, Maine, United States of America

**Keywords:** gorgonian, animal forest, canopy, planar area, frontal area

## Abstract

Octocorals have increased in abundance on many Caribbean coral reefs, and at some sites “octocoral forest” may be a better community descriptor than “coral reef.” Implicit to the concept of a forest is that structural elements, trees, colonies, etc., alter the environment in ways that affect the structural elements themselves and the organisms that inhabit the forest. At what density do the structural elements create the emergent properties of a “forest?” Communities traditionally characterized as hardgrounds and coral reefs around Puerto Rico and St John, US Virgin Islands, varied in density of octocoral colonies from a few to >100 colonies m^-2^in surveys conducted in 2021 and 2022. Canopy cover was correlated with the density of colonies. Among the quadrats with the highest octocoral density there was no significant correlation between numbers of colonies and canopy cover. Frontal area, a measure related to the community’s effect on water flow, as well as the volume occupied by colonies followed patterns similar to canopy cover. Vertical profiles of flow velocity were measured from the substratum to 3 m above the bottom on a reef on St John where octocoral population density ranged from 0 to 16 colonies m^-2^. Profiles of orbital velocity exhibited perturbations which were more pronounced in locations with > 12 colonies m^-2^. Using the effect on flow as a criterion, 4 of the 8 surveyed sites would function as forests. Understanding the density at which emergent properties appear is critical to understanding the bio-physical interactions affecting the community.

## Introduction

Animal forests are ubiquitous in the world’s oceans. Like their terrestrial analog, animal forests are defined by the arborescent organisms that form the ecosystem’s physical structure, and as in terrestrial forests, those organisms generate microhabitats that are exploited by a diversity of species (Rossi et al. 2017; Orejas et al. 2022). Recognizing an ecosystem as a forest is easy when there are hectares of trees with dense canopies that block direct sunlight, or kilometer long tracts of branching stony corals that obscure almost all views of the substratum. However, the distinction between forest and “not-forest” is less clear as the density of ecosystem engineers decreases. An operational definition of terrestrial forests used by the United Nations Food and Agriculture Organization (FAO 2018) is “an area of at least 0.5 hectares having trees higher than 5 meters and a canopy cover of more than 10 percent, or trees able to reach these thresholds *in situ*.” Implicit in the definition is the recognition that the alteration of the environment and creation of microhabitats is dependent on the size and density of trees. Creating similar criteria for marine animal forests is more difficult as the ecosystems to which the term has been applied cover organisms that span sizes that range from centimeters to 10s of meters. Orejas et al. (2022) offered a definition of a marine animal forest as a “3D structure formed by benthic animals acting as autogenic ecosystem engineers which provides new ecological niches and colonization surfaces for other organisms and results in increased provision of functions and services.” This characterization of marine forests is clear in a heuristic sense but does not provide a simple operational definition.

In this paper, we explore the nature of octocoral forests on a group of Caribbean coral reefs. We examine the question of how the number and sizes of octocoral colonies create a physical structure with emergent properties such as canopy cover and frontal area, and discuss how that structure affects conditions within the forest. We compare the physical structure of octocoral communities on reefs that were historically dominated by scleractinian corals, as well as from adjacent habitats that have been called hard grounds or gorgonian plains (sensu Williams et al. 2015). At one of the sites, we explore the relationship between density of larger octocoral colonies with the vertical structure of water movement on the reef.

Understanding Caribbean octocoral forests is particularly important as at some sites they appear to be successors to the scleractinian dominated systems that historically defined Caribbean coral reefs (Lasker et al. 2020a). The decline in abundance of scleractinians on Caribbean reefs has been well documented (Gardner et al. 2003; Jackson et al. 2014), and increasing abundances of octocorals have been reported from a variety of sites at which octocorals have been monitored (Ruzicka et al. 2013; Lenz et al. 2015; Tsounis and Edmunds 2017; Sanchez et al. 2019).

## Methods

### Abundance distributions

Octocoral abundances were determined at 6 sites on the southern shore of St John, US Virgin Islands, and 2 sites near La Parguera, Puerto Rico. The sites on St John have been monitored since 2014 and descriptions and locations are noted in Tsounis et al. (2018) and Lasker et al. (2020b). The data presented here were collected from 10 m transects arrayed parallel to each other. At three sites, Grootpan Bay, Europa Bay, and Tektite (all 7-9 m depth), ten 1 m^2^ quadrats were censused along each of 6 transects. Ten quadrats at each of four transects were monitored at Booby Rock and 10 quadrats from each of three transects at Yawzi Point (9 m depth). Quadrats were positioned randomly with respect to meter on the transect and the side of the transect. At Deep Tektite (14m) all colonies within 1 m of both sides of 3 transects were sampled, 20 quadrats on each transect. Within each quadrat the height of all upright octocorals ≥5 cm was measured. Height was measured as the distance from the base to the tip of the longest branch. Colonies were identified to species or when species could not be distinguished in the field to a species group, generally composed of 2 or 3 congeners. The St John sites were surveyed in June and July 2021.

The two transects on the south shore of Puerto Rico offshore of La Parguera were originally established by P. Yoshioka in 1983 (Yoshioka and Yoshioka 1989; Wells et al. 2022). Those sites were relocated in 2021 (Wells et al. 2022) and censused in 2022. In an effort to resurvey Yoshioka’s original transects, at Media Luna a 37 m transect was laid out and all colonies ± 0.5 m of the transect were identified to species or species group and their heights measured. At Media Luna 35 m were surveyed in the same manner.

Patterns of abundance and the physical structure of the octocoral community were analyzed using four indices. First, we characterized abundance on the basis of octocoral density (colonies m^-2^).

We then used the heights of the colonies to estimate each colony’s breadth (the widest measure of the colony parallel to the substratum) and thickness (the dimension perpendicular to breadth). In order to estimate breadth and thickness, each species was classified as either a rod, plume, bush, candelabrum or fan (after Rossi et al. 2018). Cerpovicz and Lasker (2021) had previously measured height, breadth, and thickness of 480 colonies representing 26 species at Grootpan Bay. Those colonies were assigned to the five colony forms and an average ratio between height and breadth and height and thickness calculated for each morphologic group. Heights of colonies on the quadrats were then used to estimate breadth and thickness of each colony. Using those dimensions we estimated canopy cover, the frontal area of all of the colonies within the quadrat and the volume of colonies within each quadrat. The three measures were chosen to characterize the colonies’ use of space within the quadrat (volume and canopy cover), their potential effects on water flow (frontal area and canopy cover) and the interception of downwelling light (canopy cover).

Canopy cover, frontal area and volume were all calculated by modeling each colony as an inverted cone with an ellipsoidal base (Online material Fig 1). Canopy cover was calculated as the sum of the ellipses defined by each colony’s breadth and thickness; frontal area as the sum of the triangles defined by colony height and breadth and volume as the sum of the volumes of the cones defined by each colony. As canopy cover and frontal area were scaled to 1 m^2^ areas those values are equivalent to planar area density and frontal area density, which are used in the meteorologic and hydrodynamic literature (cf. MacDonald 2000). In addition, canopy cover estimates for a subset of the quadrats were compared with values derived from analyses of images of quadrats that were acquired in 2022. Those images were analyzed using CoralNet in which the perimeter of the colonies’ canopies was defined as a convex hull and the area within the canopy quantified from the proportion of 150 randomly placed points that were within the convex hull.

Analyses were conducted using each 1 m^2^ area as a replicate. Octocoral density (the count of colonies in each m^2^), canopy cover, frontal area and volume were calculated and compared between sites using generalized linear models and pairwise comparisons made with Kruskal-Wallis Tests using Bonferroni corrected significance levels (SPSS v. 29). The relationship of octocoral density to canopy cover, frontal area and volume were explored using the regression function in Microsoft Office 365 Excel. Size frequency distributions of colony heights were compared between sites with a log linear test (SPSS v.29). The distribution of canopy cover and volume of colonies within 10 cm bands rising from the substratum were also calculated and compared between sites using log-linear tests (SPSS v.29).

### Flow effects

A high-resolution acoustic Doppler current profiler (Nortek Aquadopp 2 MHz) was laid on the bottom, measuring upwards, in 85 locations spread among three sites with depth ranging from 5.7 to 13.2 m (Grootpan Bay and Europa Bay) at St John USVI in March 2016, March and August 2017, July and November 2018 and March and November 2019. The 85 locations were selected to cover the range of substratum composition ranging from bare seabed (no octocorals) to the densest population of octocorals where the acoustic beams of the current profiler could travel without reflection on octocoral branches. Population density of octocorals was determined from counts of individuals > 5 cm tall in 4 plots of 1m^2^ area surrounding the current profiler.

The current profiler was set to Pulse Coherent Mode, enabling measurement of high-resolution high-frequency flow velocity profiles. Flow velocity was measured every 3 cm from 0.3 to 2.3 m above the seabed for a total of 68 simultaneous time series of 5 min at 2Hz sampling frequency repeated every 15 min during 2-12h. A quality check on the velocity data was applied following Nortek recommendations, removing data with less than 70% correlation between acoustic beams. Ambiguous velocity determinations, inherent to the Pulse Coherent Mode, were detected using the methodology of Goring and Nikora (2002) and corrected following Nortek recommendations (https://support.nortekgroup.com/hc/en-us/articles/360013106500-TN-028-Aquadopp-HR-Profiler-Extended-Velocity-Range-Mode-2010). Less than 3% of the data required correction. To get reliable velocity statistics per burst (mean, U_c_ and standard deviation, U_rms_), a threshold of 60% valid data per burst was applied. In total, 455 velocity profiles were validated, meaning an average replication of 4 bursts per location. Values at 2 m above the seabed were taken as a reference to characterize the flow conditions at each location and time and to normalize the values along the vertical profiles.

## RESULTS

### Abundance distributions

The sites differed markedly in appearance (Fig. 1). At the two sites from Puerto Rico, and to a lesser extent at Booby Rock and Grootpan Bay on St John, octocorals dominated the seascape. The four other St John sites had fewer octocoral colonies, but octocorals were still the visually most common macrobenthic invertebrates. That subjective characterization of sites by octocoral prominence was supported by all measures of octocoral abundance (Fig. 2 and GLM comparisons, p<0.001 for all measures of abundance [Online Table 1]). Pairwise comparisons indicated that the sites could be divided into three groups, Media Luna and San Cristobal which had dramatically greater abundances than the St John sites regardless of measure, and then a second group composed of Booby Rock and Grootpan followed by a third group composed of the other four St John sites.

**Fig 1.**
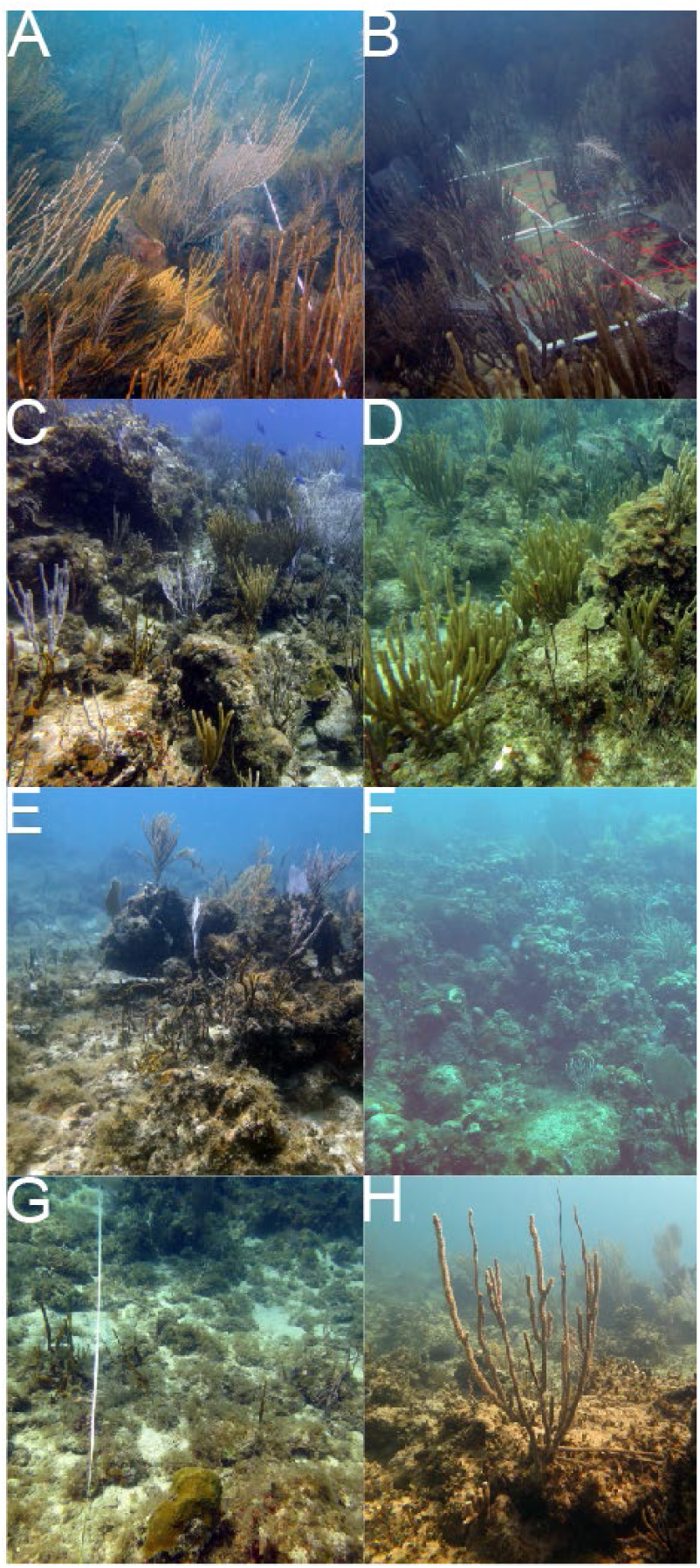
Images of the communities at each of the study sites. A, Media Luna; B, Cristobal; C, Booby Rock; D, Grootpan; E, Yawzi; F, Deep Tektite; G, Tektite; H, Europa

**Fig 2.**
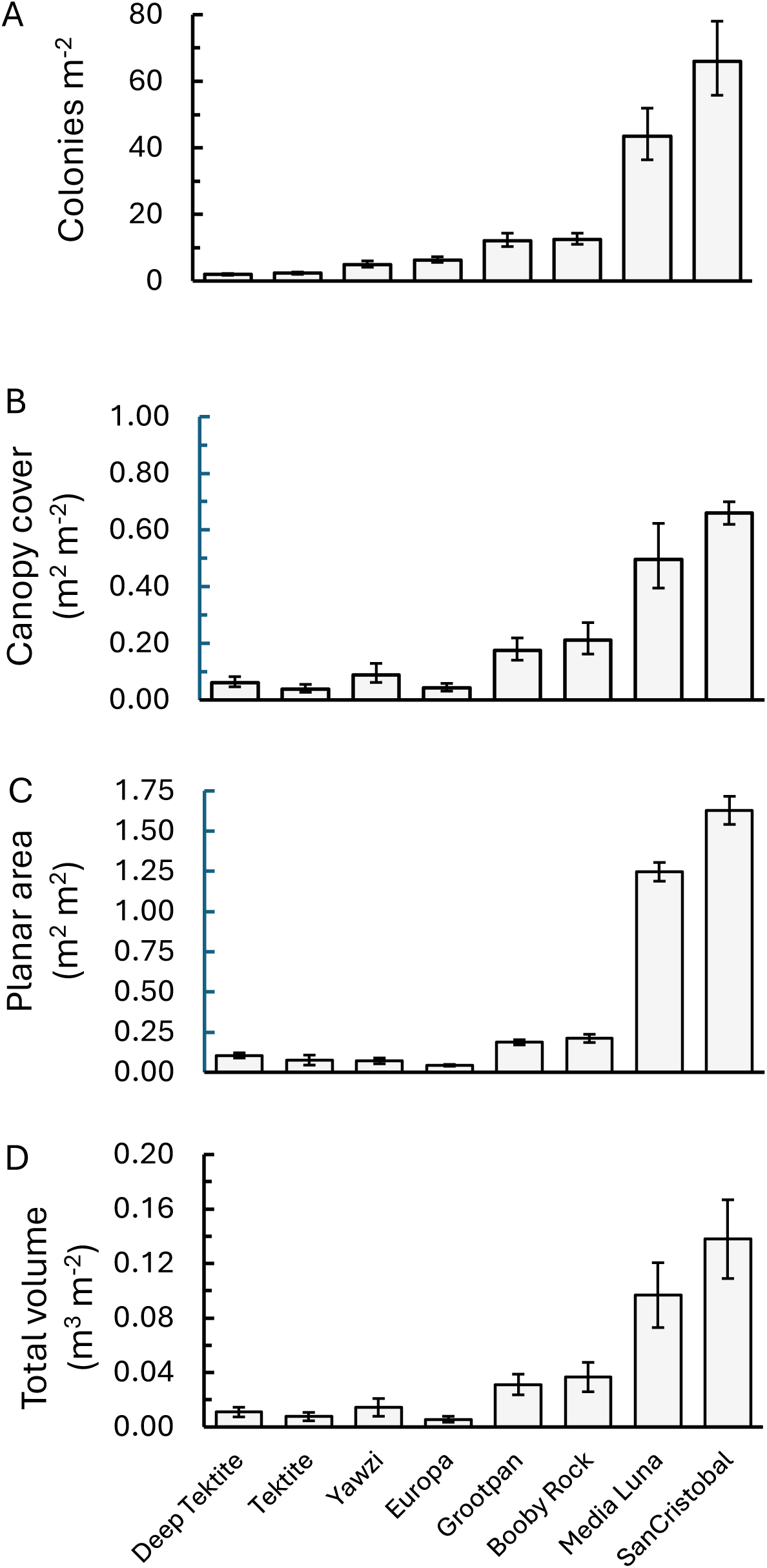
Average density (A), summed canopy cover (B), summed planar area (C) and summed volume (D) (+/-standard error) of ≥ 5 cm tall octocoral colonies found in 1 m^2^ areas at study sites from Puerto Rico and St John

Size frequency distributions of colonies at the sites were significantly different from each other (Online Fig. 2, Log-linear test, p<0.001). However, the general trend was for the greatest proportion of colonies to be in the 5-10 and 11-20 cm size classes. The proportion of small colonies was greatest at Europa Bay and Yawzi Point, two of the sites with low octocoral density. Large colonies were relatively more common at Deep Tektite, which also had low octocoral density.

Like the differences in octocoral density, the estimated canopy cover at the Puerto Rico sites was greater than at the St John sites (Online Table 1). When considered as either individual 1m^2^ quadrats or quadrats averaged by site (Fig. 3), canopy cover increased with increasing density of colonies. Two quadrats at San Cristobal had octocoral cover > 100%, an outcome resulting from the methodology which did not correct for colonies with overlapping canopies. Canopy cover was linearly related to octocoral density with r^2^ values of 0.81 using each m^2^ and 0.98 comparing site averages. However, the relationship between octocoral density and cover among the high-density Puerto Rico quadrats was not as clear. Canopy cover at those two sites was significantly correlated with octocoral density (p<0.01), but the best fit model, a logarithmic function only explained 18% of the variance. Canopy cover among the high-density quadrats averaged approximately 60% (5829 cm^2^, SE = 217 cm^2^). The comparison of canopy cover measured as ellipses versus that from point counts of the area within canopies showed strong correlation between the two measures (see Online Fig. 3 and associated discussion). Although the 8 sites differed dramatically in overall canopy cover, the height of the canopy was similar across most of the sites (Online Fig. 4). Canopy cover peaked between 30 and 50 cm for all sites with the distribution displaced to somewhat higher heights at Grootpan and Booby Rock.

**Fig 3.**
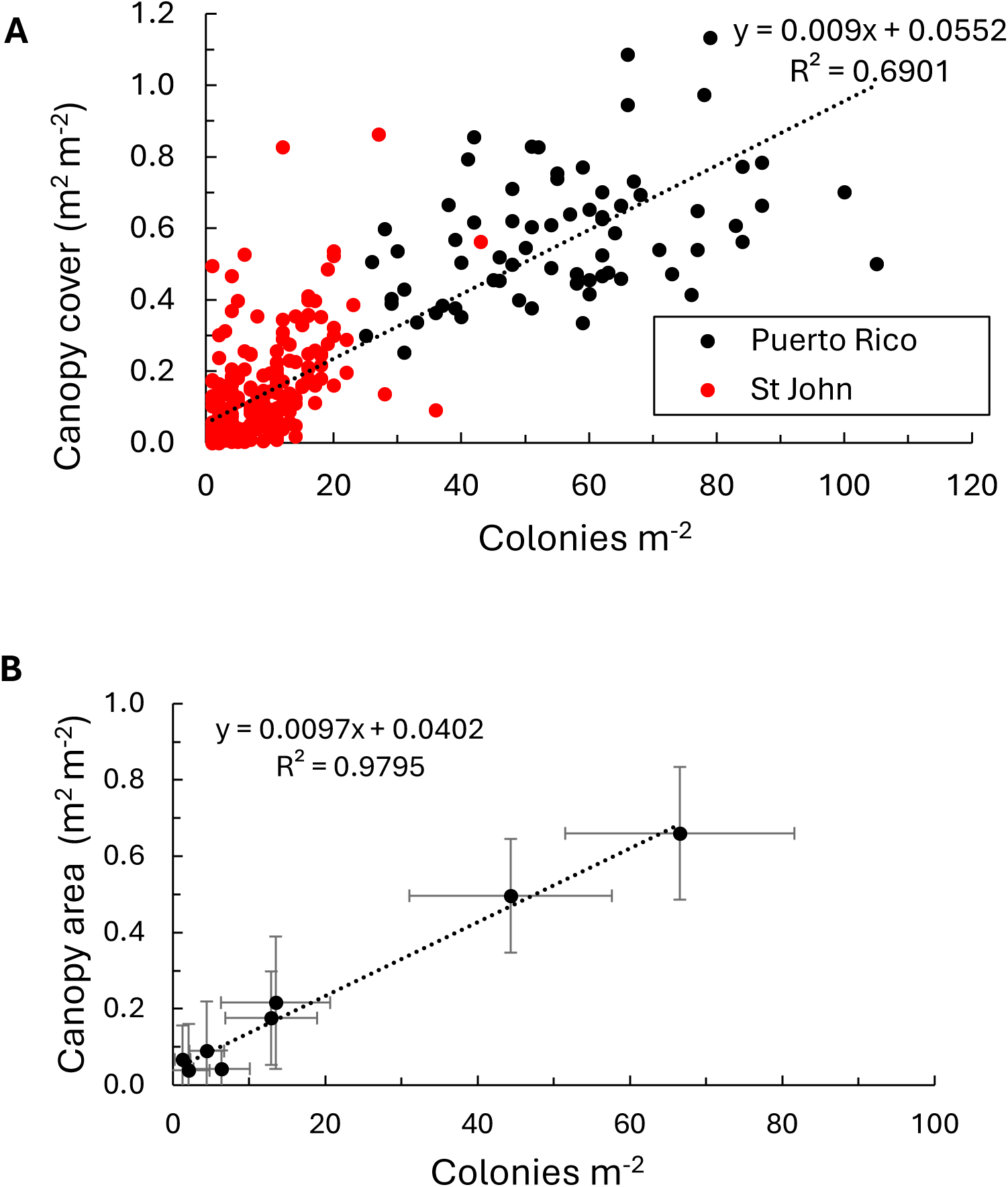
Summed canopy cover of octocoral colonies in 1 m^2^ quadrats as a function of octocoral density. A, all colonies; B, averages at each site with standard deviation

Frontal area ranged from near zero to almost 3.0 m^2^ m^-2^ (Fig. 4A). Frontal area represents the area occupied by colonies when viewed sideways. Values greater than 1.0 are possible when the frontal area of individual colonies that are behind each other in the quadrat are summed. Frontal area increased with octocoral density, and a linear model best described the relationship, explaining 86% of the variance in frontal area (Fig. 4A). A logarithmic relationship was the best ft to the data when frontal area was analyzed separately for the Puerto Rico sites (r^2^=0.18). Frontal area at the St John sites best fit a linear model (r^2^=0.49).

**Fig 4.**
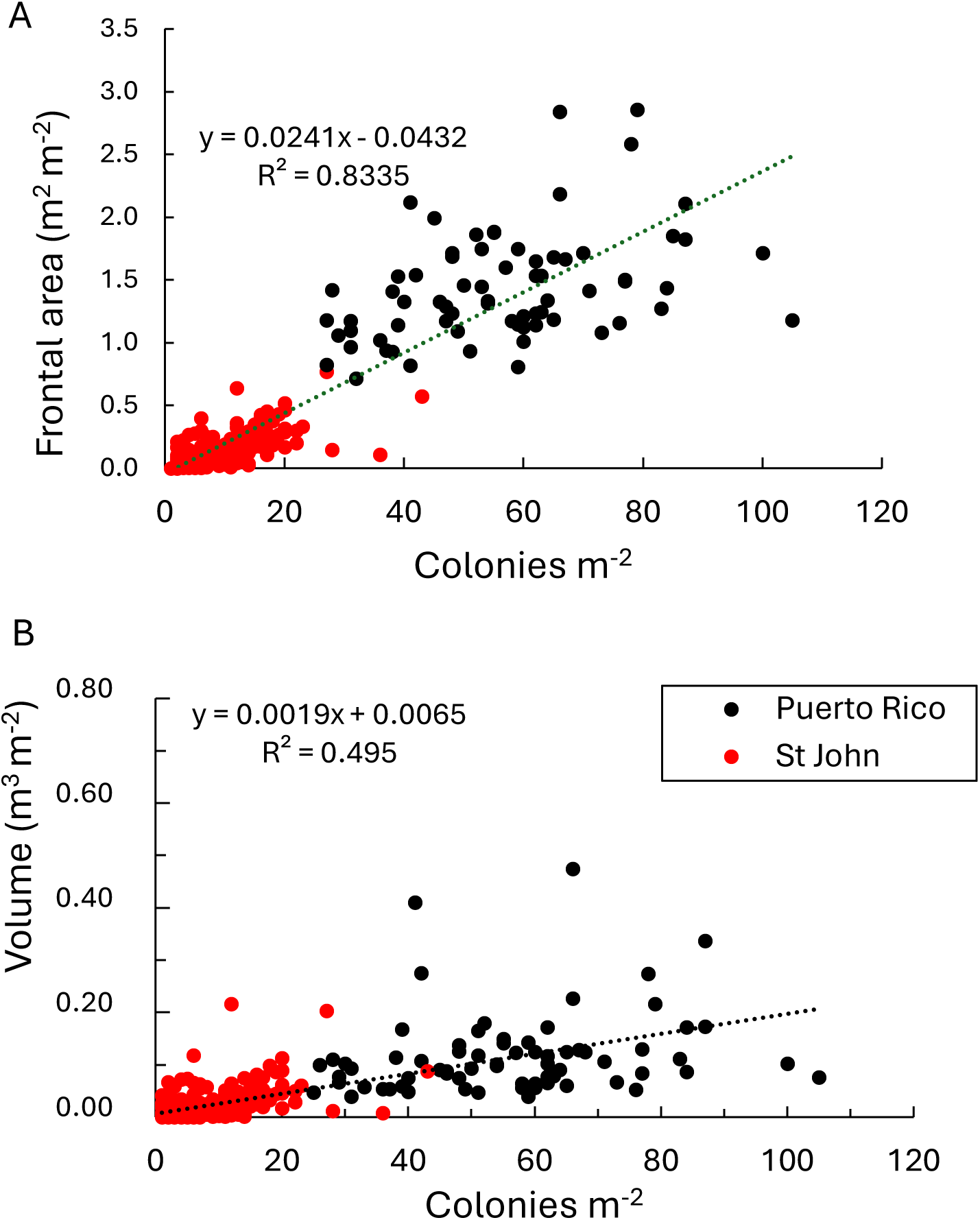
A, Summed frontal areas of colonies within each 1 m^2^ quadrat at sites on Puerto Rico and St John as a function of octocoral density; B, Summed canopy volume of colonies within 1 m^2^ areas at sites on Puerto Rico and St John in 2021 as a function of octocoral density

Mirroring canopy cover and frontal area, the estimated volume occupied by colonies at the Puerto Rico sites was greater than the St John sites (Fig 4B). However, in most cases, octocorals occupied less than 20% of the available volume in the cubic meter above the quadrat. The greatest volume observed was less than 50% of the available space. There was a significant relationship between octocoral density and canopy volume (Fig. 5), but there was no significant relationship between volume and octocoral density among the Puerto Rico m^2^ quadrats (r^2^=0.05, p>0.05). Volume at the Puerto Rico sites averaged 0.118 m^3^ (std error = 0.010 m^3^) which corresponds to approximately 12% of the volume of the cubic meter immediately above the substratum. When the volume occupied by colonies in the water column above the substratum is divided into 10 cm thick layers it becomes apparent that most of the canopy was found in the layers between 30 and 50 cm (Online Fig. 5). At Media Luna and San Cristobal, the sites with the greatest octocoral density, the colonies occupied 15-20% of the volume in the band 40 cm above the substratum. Those values range from 1 to 7% at the St John sites.

**Fig 5.**
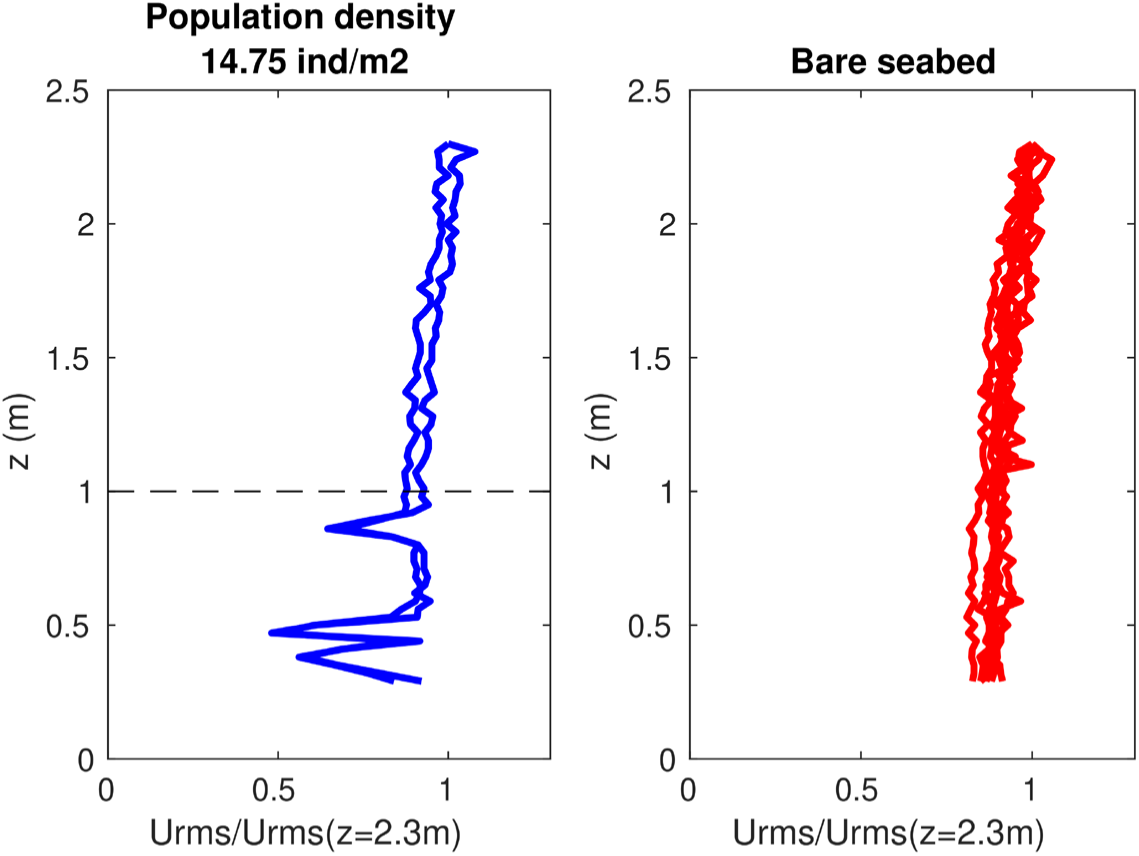
Vertical profile of wave orbital velocity (standard deviation of 5 min flow velocity record) above a bare seabed (right) and in a location with octocoral population density of 14.75 individuals.m^-2^ (left) on a fringing reef in Grootpan Bay, St John, US Virgin Islands. Standard deviation is normalized to its value at 2.3 m above the bed

### Flow effects

Current was profiled at 85 locations, and octocoral density ranged from 0 to 16 individuals m^-2^ at those locations. Wave action dominated the flow during most observations, indicated by 90% of the dataset showing a U_rms_ to U_c_ ratio > 2 at 2.3 m above the seabed. In this group of flow profiles, U_c_ at 2.3 m above the seabed was < 4.6 cm/s while U_rms_ at 2.3 m ranged from 3 to 16 cm/s. Vertical profiles of U_rms_ normalized by their value at 2.3 m above the seabed displayed a smooth decay from 2.3 to 0.3m on a bare seabed (Fig. 5 and Online Fig. 5). In the presence of high octocoral densities, vertical profiles of U_rms_ displayed fluctuations up to 1 m above the seabed.

The fluctuations of wave orbital motion were quantified using the standard deviation of normalized U_rms_ along two separate portions of the vertical profile, up to 1 m above the seabed and in the 1–2.3 m layer (Fig. 5). The statistics (median and 10% and 90% quantiles) of standard deviations of normalized U_rms_ in the two different depth layers were calculated across replicates for each of the 85 locations. In four out of 15 locations where octocoral population density was larger than approximately 10 individuals m^-2^, wave orbital motion was clearly more perturbed near the bed than above it, while that was not the case at lower density (Fig. 6).

**Fig 6.**
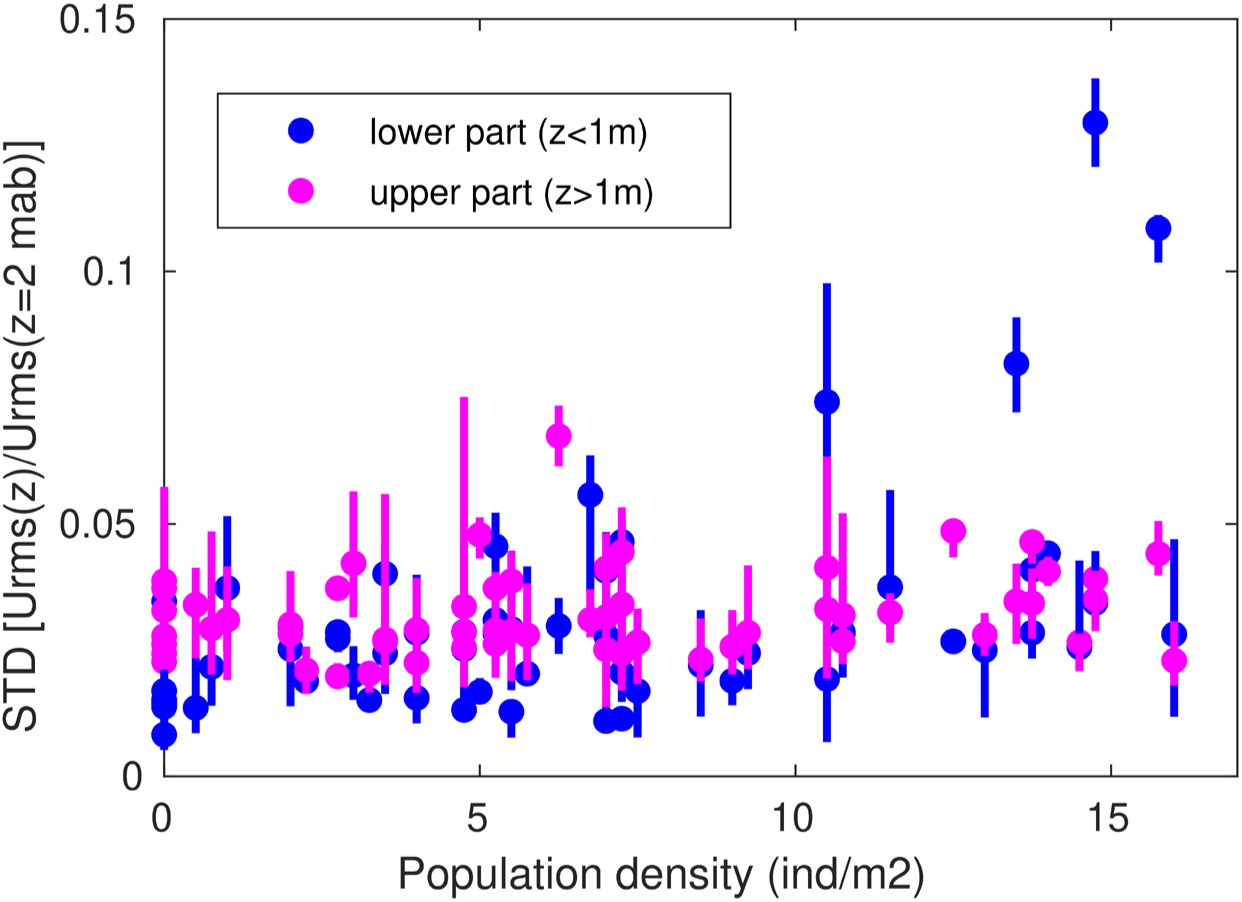
Standard deviation of orbital velocity fluctuations across the lower (from 0.3 to 1 m above the seabed) and the upper (from 1 to 2.3 m above the seabed) part of the vertical profile t vs octocorals population density at different locations on a fringing reef in Grootpan Bay, St John, US Virgin Islands

In 12 flow profiles, 3% of the dataset, wave motion did not dominate the flow (U_rms_ /U_c_ <1). The Octocoral densities associated with those measurement ranged from 0 to 10.5 individual per meter square. In these latter flow profiles, the mean flow (normalized by its value at 2.3 m above the bed) decreased near the bed more rapidly in the presence of octocorals than on a bare seabed, and the mean flow attenuation was larger at the largest octocoral population density (Online Fig. S6).

## Discussion

If the definition of an animal forest incorporates the criterion that the structural elements of the forest alter the physical and biotic environment, then the relevant question becomes at what combination of colony density and colony sizes do the octocorals produce such an effect? The sites surveyed for this study were selected because gorgonian octocorals were visually the most common macro-invertebrates in the benthic community. Abundances at the sites, as well as the related traits of canopy cover, the volume occupied by the colonies and the frontal area varied considerably, illustrating the gradient from octocoral dominated communities to octocoral forests.

The physical structure created by forests is an emergent property of the ecosystem in that the compounded effect of multiple individuals is fundamentally different than that of a single individual. In the octocoral forests these effects fall into multiple categories. The trait that is immediately visible is that of octocoral branches intercepting downwelling light and thus altering light levels on the substratum (Girard and Edmunds 2023). Octocorals at the study sites did not form the dense layer of light absorption found in leafy terrestrial forests or underwater forests such as kelp beds. Our estimates of the canopy cover in the densest populations were generally at 60%, and canopy cover at the low octocoral density sites was 4%. Those characterizations treated the canopy created by each colony as a solid ellipsoidal disc. That approach overestimates the effects of colonies, as light interception is a function of the number and size of the branches within the perimeter of the ellipse. Girard and Edmunds (2023) measured “canopy closure” at sites on the south shore of St John based on the proportion of upward directed images that were occupied by branches. They reported a positive relationship between canopy closure and colony density, and report mean closure (i.e., canopy cover) at Grootpan Bay (labeled East Cabritte in their study) of 31.4%. In our analyses quadrats at Grootpan Bay in the upper quartile of canopy cover values ranged from 26 to 49% cover.

Girard and Edmunds (2023) also found the canopy cover at Grootpan Bay caused a 33% reduction in light intensity below the canopy. The St John sites both in our survey and that of Girard and Edmunds (2023) had lower canopy cover than the Puerto Rico sites. Thus, it is likely that the canopy at sites with densities like those of Grootpan and Booby Rock on St John as well as the greater octocoral density sites at Media Luna and San Cristobal substantively reduce light levels at the substratum. Data from the Puerto Rico sites suggest canopy cover plateaus at high colony densities. That in conjunction with the diffraction of light in the water column and reflection of light from the substratum should set an upper limit on the effects of the canopy on light levels. All but the lowest octocoral density sites in this study (Tektite, Deep Tektite, Europa Bay and Yawzi) are likely to affect light levels and even at sites with low average octocoral densities, there should be light effects within those quadrats that contain multiple colonies.

In terrestrial forests, another emergent property of canopies relates to wind attenuation within the forest, and uplift of the turbulence peak at the top of the forest with significant alteration of the flow just above the canopy (Raupach et al. 1996). In aquatic systems, similar effects on steady currents within and above marine animal forests are expected (Monismith 2007; Nepf 2012), with consequence for the transport and diffusion between the overlying flow and inside the canopy (Falter et al. 2007).

Flow transport attenuation has been related to planar and frontal area density of obstacles, with a linear increase of the flow attenuation coefficient from 0.5 to 3 when frontal area density increased from 5 to 35% (Macdonald 2000). Flow transport attenuation in the canopy may have the greatest effects on the aquatic community. The frontal areas reported in Fig. 5 ranged from near zero to almost 3 m^2^ m^-2^, i.e. from less than 1% to almost 300% of a 1 m^2^ “wall” bounding the quadrat. Those values are overestimates because we treated each colony’s frontal area as a solid. However, even with corrections for those effects McDonalds’s work suggests the quadrats with large frontal areas would have strong effects on flow transport.

In the present study, flow transport rates near the bed were already negligible outside octocoral populations most of the time, preventing a test of the effect of the canopy’s frontal area on flow attenuation. In our observations, the flow around the octocorals was dominated by wave orbital motion. Lowe et al. (2005)established theoretically and experimentally that wave orbital motion decreased when either planar or frontal area density increased above and within a canopy formed by stiff obstacles. Similarly, Abdolahpour et al. (2017) measured a two-fold reduction of orbital velocity inside a flume with a dense canopy of artificial stiff stems with 10% planar area density. No significant reduction was observed inside a sparse canopy with 1% planar area density. In the present field study, the density of octocorals ranged from 1-16 colonies m^2^, which based on the regression in Fig. 4 would have frontal areas of 0-42%. At those frontal areas, wave orbital velocities were not consistently lower in the canopy compared to above it, in contradiction with the findings of Lowe et al. (2005) and Abdolahpour et al. (2017). Our discrepant observations are likely due to octocorals’ flexibility. In aquatic vegetation composed of flexible organisms, a coherent waving (the monami) has been observed in response to vortex passage at the top of the canopy moving up and down flexible vegetation stems (Ghisalberti and Nepf 2006). Similarly, oscillation of the flexible octocorals under wave oscillatory motion could likely explain why a reduction of wave orbital motion was not observed. However, we did observe an increase in the variability of velocity within the canopy at densities larger than 10 colonies m^-2^. Based on the regressions in Figs 3 and 4 a quadrat with 10 colonies m^-2^ would have a canopy cover of 15 % and frontal area of 20 %. Given the commonness of octocorals in shallow tropical waters (c.f., Reichelt et al. 1986; Norstrom et al. 2009; Yranzo et al. 2014; Velásquez and Sánchez 2015; Edmunds et al. 2016; Benayahu et al. 2019; Kupfner Johnson and Hallock 2020; Lasker et al. 2020a; Lalas et al. 2024; Otis et al. 2024) where wave driven flow often dominates, the present findings likely apply to many localities.

Effects of octocorals on physical processes such as sedimentation and flow measured as the dissolution of clod cards were examined by Cerpovicz and Lasker (2021). Working in Grootpan Bay they compared sedimentation and dissolution of clod cards in paired canopy and non-canopy locations, using an octocoral density of 12 or more colonies within a meter of the sediment and clod card to define the canopy locations (i.e., ≥4 colonies m^-2^). Sediment traps deployed within the octocoral canopy accumulated greater amounts of sediment, with coarser grains and higher organic content than paired traps placed outside the canopy. Clod cards had greater rates of dissolution within the canopy sites. These results suggest greater turbulence and resuspension within the canopy.

Other environmental and ecological effects of the marine animal forest also have been observed (Nelson and Bramanti 2020; Bosch et al. 2023). For example, hydrodynamic effects may affect the supply of particulate food, and the supply and settlement of larvae. Privitera-Johnson et al. (2015) compared the octocoral density of colonies with recruitment at sites on St John, including 4 of the sites in this study. They found a positive correlation between adult density and recruitment in both 2013 and 2014 for all octocorals, for *Gorgonia* spp. and for plexaurid species but not among *Antillogorgia* spp. They suggested the relationship was generated, in part, by the eddies created by adult colonies, which would entrain passing larvae. However, net flow at sites with wave driven oscillations will be low and the relationship they observed may be related to some other feature of the habitat that correlates with adult densities. In other systems, the density of seaweeds differentially affects the recruitment of seaweeds and microphytobenthic organisms (Umanzor et al. 2018).

The effects of octocorals forests may also extend to the settlement and survival of other taxa. Girard and Edmunds (2023) found a correlation between benthic invertebrate community structure and canopy metrics at two of those sites. The site where there was no correlation also had the lowest octocoral density and canopy closure (4.1 colonies m^-2^ and 7.9% closure, respectively). The canopy also is critical in considering interactions among octocoral colonies. Colonies that grow into each other suffer damage as branches abrade, leaving sharp lines of demarcation where either direct interactions or altered growth patterns keep colonies from interdigitating (Gambrel and Lasker 2016). Thus, the development of an octocoral forest may differentially affect the species making up the forest.

When do octocorals become an octocoral forest, kelp a kelp forest, corals a coral reef, or blades of seagrass a meadow? Falter et al. (2007) modeled flow on a coral reef with rigid structures and concluded that “A true coral canopy is developed when the spacing of the individual colonies is on the order of the colony height or closer and the horizontal expanse of the aggregation is greater than roughly 10 times the canopy height.” Following this logic, 40 cm tall octocoral colonies require a density of 7 evenly spaced colonies m^-2^ and 50 cm colonies require of 5 colonies m^-2^. In the case of octocorals, our and other empirical studies suggest emergent properties become apparent at densities around 5-10 colonies m^-2^, which corresponds to ∼ 10% canopy cover. The FAO threshold of 10% for terrestrial forests is surprisingly suitable for these marine animal forests. However, more precise estimates of frontal area and canopy cover based on measures of the actual colonies, like those from point counts of images, are necessary to ascertain this threshold. Similarly, it is important to ascertain whether effects of the physical structure steadily increase as colony density increases, or there may be thresholds below which the effects of the octocorals are negligible. Understanding the mechanistic relationship between species individual shape and spatial density and flow perturbation is essential to anticipate changes in the emergent forest effect associated with changing octocoral communities under the effects of climate change.

## Acknowledgements

The research was conducted with support from the U.S. National Science Foundation (HRL, OCE 1756381; PJE, OCE 13-32915 and OCE 17-56678). LB was funded by CNRS (PICS project GORGOSHIFT) and from BNP PARIBAS Foundation (project DEEPLIFE). The French Ministère de l’Enseignement Supérieur et de la Recherche funded NP through a doctorate contract. We thank the staff of the University of Puerto Rico Isla Magueyes Marine Laboratory and the University of the Virgin Islands Environmental Resource Station for their assistance facilitating out research. E.A. Anderson and J. Benz assisted with fieldwork on St. John. The research was conducted under research permits from the U.S. National Park Service (VIIS-2021-SCI-0008) and the Commonwealth of Puerto Rico Department of Natural and Environmental Resources.

## Online Material

**Online Fig 1.**
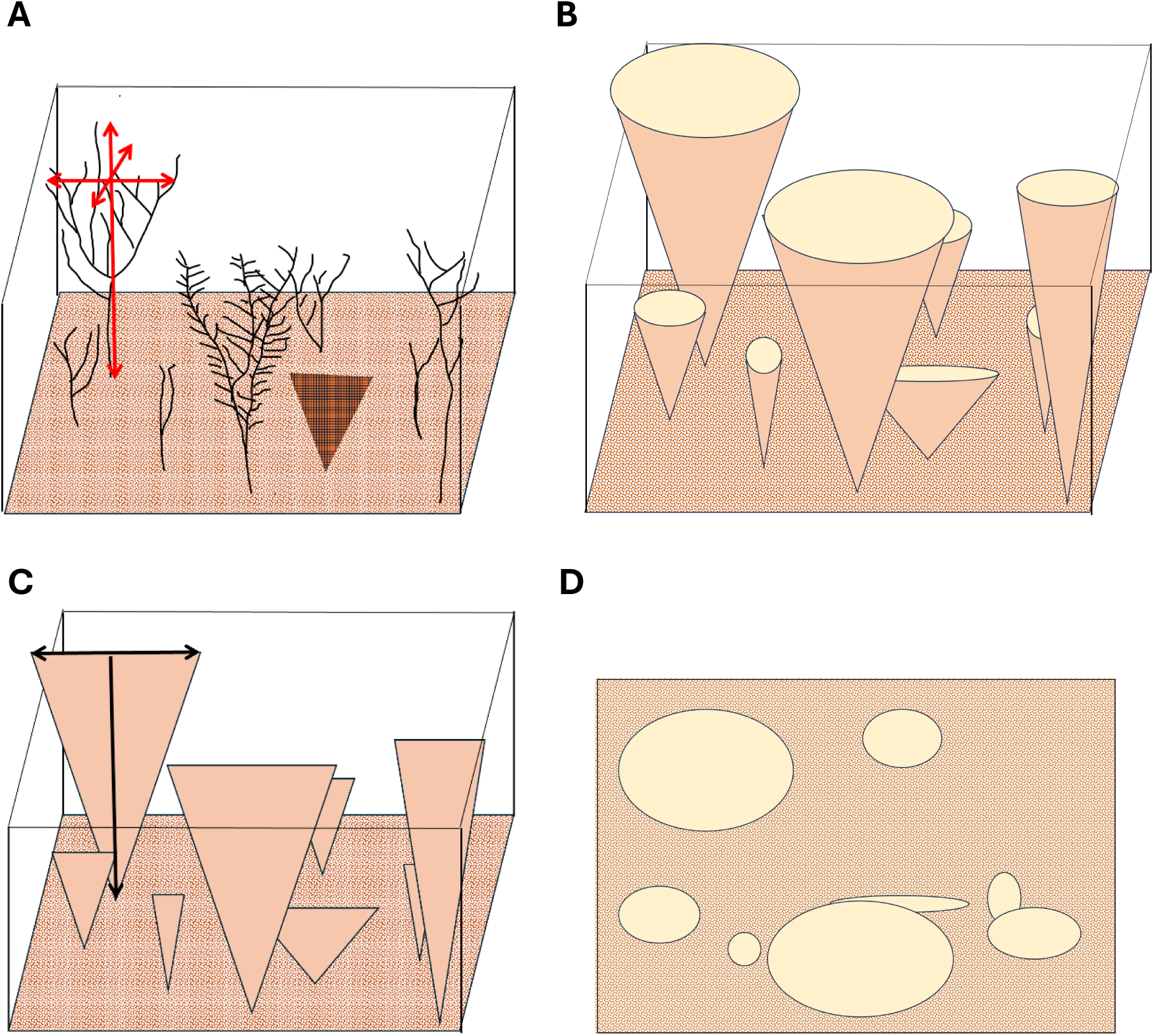
A, depiction of colonies on the substratum each with a height, breadth and thickness. B, inverted ellipsoidal pyramid models of the colonies. Volumes of the cones were summed to calculate octocoral volume. C, Triangles used to calculate frontal area. D, ellipses used to calculate canopy area.

**Online Table 1** – Comparisons of density, canopy area and summed colony volumes of octocorals across 8 sites on Puerto Rico and St John. All analyses conducted using SPSS (v.29)

## Density

Generalized Linear Models using Ln(Sqrt) transformed values of density

Tests of Model Effects (Type III)

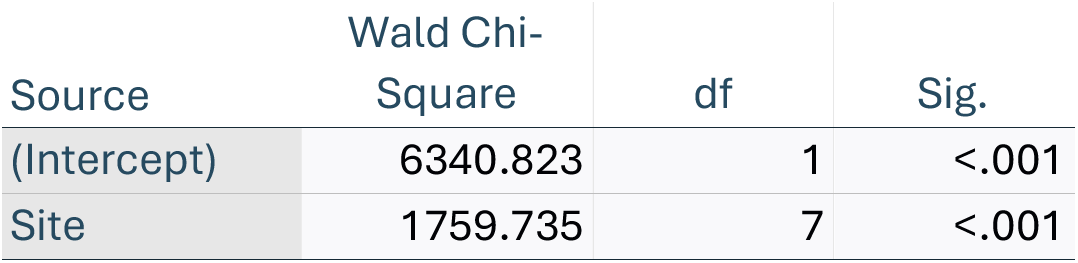

Estimated Marginal Means

| Site | Mean | Std. Error | 95% Wald Confidence Interval |  |
| --- | --- | --- | --- | --- |
|  |  |  | Lower | Upper |
| Booby Rock | 1.2501 | .04171 | 1.1684 | 1.3319 |
| Deep Tektite | .3382 | .03391 | .2717 | .4046 |
| Europa | .9253 | .03362 | .8594 | .9912 |
| Grootpan | 1.2657 | .03391 | 1.1992 | 1.3321 |
| Media Luna | 1.8862 | .04534 | 1.7974 | 1.9751 |
| San Cristobal | 2.0947 | .04282 | 2.0108 | 2.1786 |
| Tektite | .4215 | .03759 | .3479 | .4952 |
| Yawzi | .8036 | .04755 | .7104 | .8968 |

Independent-Samples Kruskal-Wallis Test

|  |  |
| --- | --- |
| Total N | 365 |
| Test Statistic | 296.337<sup>a</sup> |
| Degree Of Freedom | 7 |
| Asymptotic Sig.(2-sided test) | <.001 |

a. The test statistic is adjusted for ties.

Pairwise Comparisons of Sites

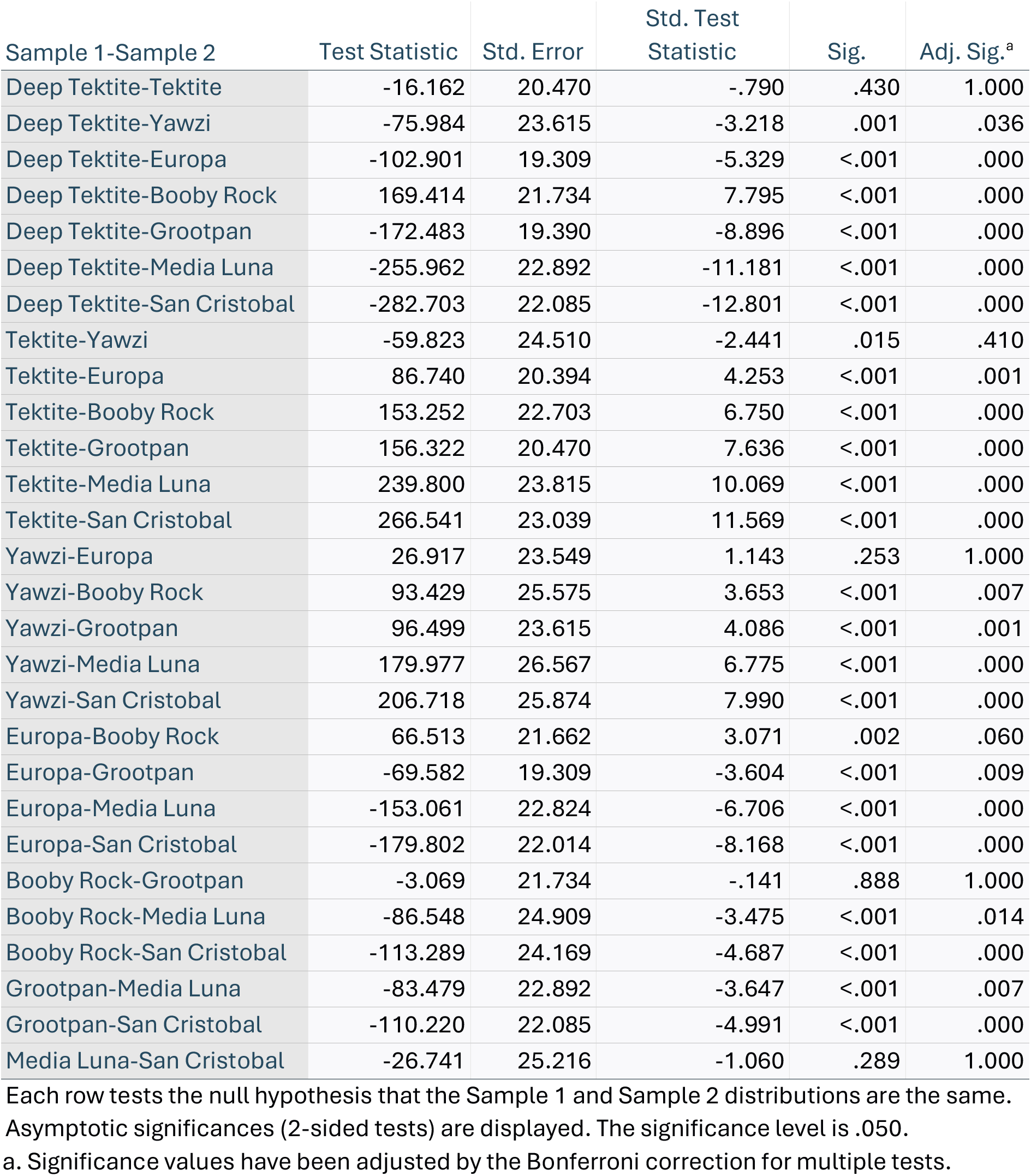

## Canopy Area

Generalized Linear Models-Canopy Area using TWEEDIE distribution

Tests of Model Effects (Type III)

| Source | Wald Chi-Square | df | Sig. |
| --- | --- | --- | --- |
| (Intercept) | 398.608 | 1 | <.001 |
| Site | 262.054 | 7 | <.001 |

Dependent Variable: Area
Model: (Intercept), Site

Estimated Marginal Means:

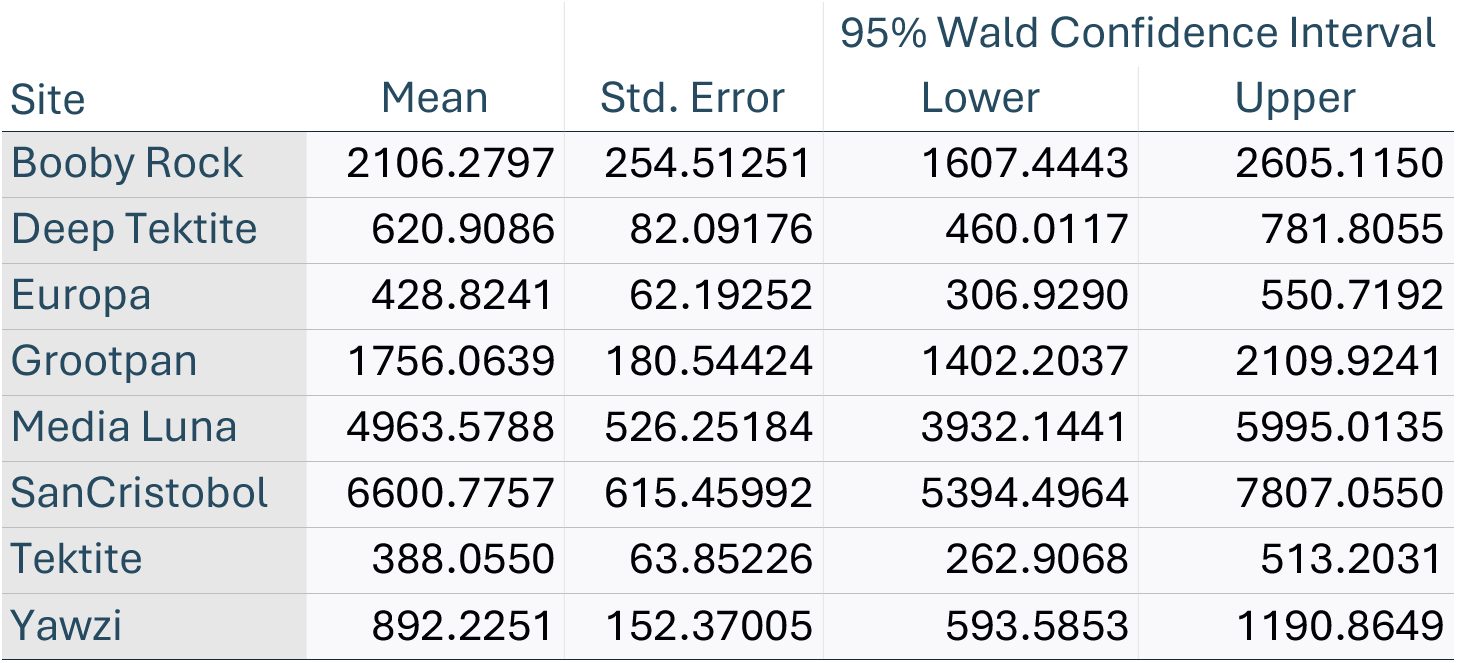

| Site | Mean | Std. Error | 95% Wald Confidence Interval |  |
| --- | --- | --- | --- | --- |
|  |  |  | Lower | Upper |
| Booby Rock | 2106.2797 | 254.51251 | 1607.4443 | 2605.1150 |
| Deep Tektite | 620.9086 | 82.09176 | 460.0117 | 781.8055 |
| Europa | 428.8241 | 62.19252 | 306.9290 | 550.7192 |
| Grootpan | 1756.0639 | 180.54424 | 1402.2037 | 2109.9241 |
| Media Luna | 4963.5788 | 526.25184 | 3932.1441 | 5995.0135 |
| SanCristobol | 6600.7757 | 615.45992 | 5394.4964 | 7807.0550 |
| Tektite | 388.0550 | 63.85226 | 262.9068 | 513.2031 |
| Yawzi | 892.2251 | 152.37005 | 593.5853 | 1190.8649 |

Independent-Samples Kruskal-Wallis Test Summary

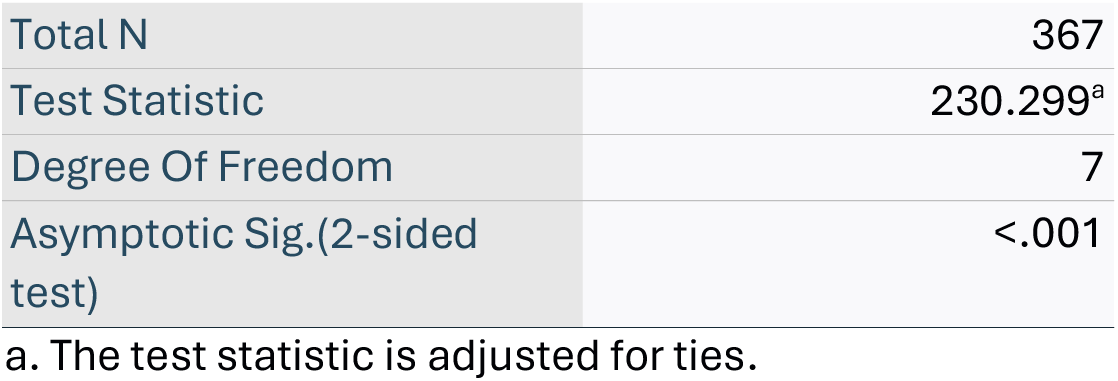

|  |  |
| --- | --- |
| Total N | 367 |
| Test Statistic | 230.299<sup>a</sup> |
| Degree Of Freedom | 7 |
| Asymptotic Sig.(2-sided test) | <.001 |

a. The test statistic is adjusted for ties.

Pairwise Comparisons of Sites

| Sample 1-Sample 2 | Test Statistic | Std. Error | Std. Test<br>Statistic | Sig. | Adj. Sig.<sup>a</sup> |
| --- | --- | --- | --- | --- | --- |
| Tektite-Deep Tektite | 34.486 | 20.417 | 1.689 | .091 | 1.000 |
| Tektite-Europa | 40.952 | 20.417 | 2.006 | .045 | 1.000 |
| Tektite-Yawzi | -64.619 | 24.582 | -2.629 | .009 | .240 |
| Tektite-Grootpan | 133.125 | 20.495 | 6.495 | <.001 | .000 |
| Tektite-Booby Rock | 143.875 | 22.755 | 6.323 | <.001 | .000 |
| Tektite-Media Luna | 230.755 | 23.879 | 9.664 | <.001 | .000 |
| Tektite-San Cristobal | 255.218 | 23.095 | 11.051 | <.001 | .000 |
| Deep Tektite-Europa | -6.467 | 19.360 | -.334 | .738 | 1.000 |
| Deep Tektite-Yawzi | -30.133 | 23.711 | -1.271 | .204 | 1.000 |
| Deep Tektite-Grootpan | -98.639 | 19.442 | -5.074 | <.001 | .000 |
| Deep Tektite-Booby Rock | 109.390 | 21.811 | 5.015 | <.001 | .000 |
| Deep Tektite-Media Luna | -196.270 | 22.981 | -8.540 | <.001 | .000 |
| Deep Tektite-San Cristobal | -220.732 | 22.165 | -9.959 | <.001 | .000 |
| Europa-Yawzi | -23.667 | 23.711 | -.998 | .318 | 1.000 |
| Europa-Grootpan | -92.172 | 19.442 | -4.741 | <.001 | .000 |
| Europa-Booby Rock | 102.923 | 21.811 | 4.719 | <.001 | .000 |
| Europa-Media Luna | -189.803 | 22.981 | -8.259 | <.001 | .000 |
| Europa-San Cristobal | -214.266 | 22.165 | -9.667 | <.001 | .000 |
| Yawzi-Grootpan | 68.506 | 23.778 | 2.881 | .004 | .111 |
| Yawzi-Booby Rock | 79.256 | 25.751 | 3.078 | .002 | .058 |
| Yawzi-Media Luna | 166.136 | 26.749 | 6.211 | <.001 | .000 |
| Yawzi-San Cristobal | 190.599 | 26.052 | 7.316 | <.001 | .000 |
| Grootpan-Booby Rock | 10.751 | 21.883 | .491 | .623 | 1.000 |
| Grootpan-Media Luna | -97.631 | 23.050 | -4.236 | <.001 | .001 |
| Grootpan-San Cristobal | -122.093 | 22.237 | -5.491 | <.001 | .000 |
| Booby Rock-Media Luna | -86.880 | 25.081 | -3.464 | <.001 | .015 |
| Booby Rock-San Cristobal | -111.343 | 24.335 | -4.575 | <.001 | .000 |
| Media Luna-San Cristobal | -24.463 | 25.389 | -.964 | .335 | 1.000 |

Each row tests the null hypothesis that the Sample 1 and Sample 2 distributions are the same. Asymptotic significances (2-sided tests) are displayed. The significance level is .050.
a. Significance values have been adjusted by the Bonferroni correction for multiple tests.

## VOLUME

Generalized Linear model using Tweedie-log linked distribution Tests of Model Effects (Type III)

| Source | Wald Chi-Square | df | Sig. |
| --- | --- | --- | --- |
| (Intercept) | 38789.745 | 1 | <.001 |
| Site | 562.717 | 7 | <.001 |

Dependent Variable: volume
Model: (Intercept), Site

Estimated Marginal Means

| Site | Mean | Std. Error | 95% Wald Confidence Interval |  |
| --- | --- | --- | --- | --- |
|  |  |  | Lower | Upper |
| Booby Rock | 36587.8600 | 4818.99346 | 28263.4455 | 47364.0590 |
| Deep Tektite | 10905.1300 | 1567.23238 | 8228.1139 | 14453.1132 |
| Europa | 5651.4238 | 957.25726 | 4054.8845 | 7876.5725 |
| Grootpan | 31201.2547 | 3476.87376 | 25079.5335 | 38817.2411 |
| Media Luna | 96808.0394 | 10868.32164 | 77687.2618 | 120634.9185 |
| San Cristobal | 137880.3181 | 13381.71356 | 113996.2383 | 166768.5040 |
| Tektite | 7557.2631 | 1317.22889 | 5370.3222 | 10634.7856 |
| Yawzi | 14357.0928 | 2724.11312 | 9898.2775 | 20824.4428 |

Independent-Samples Kruskal-Wallis Test

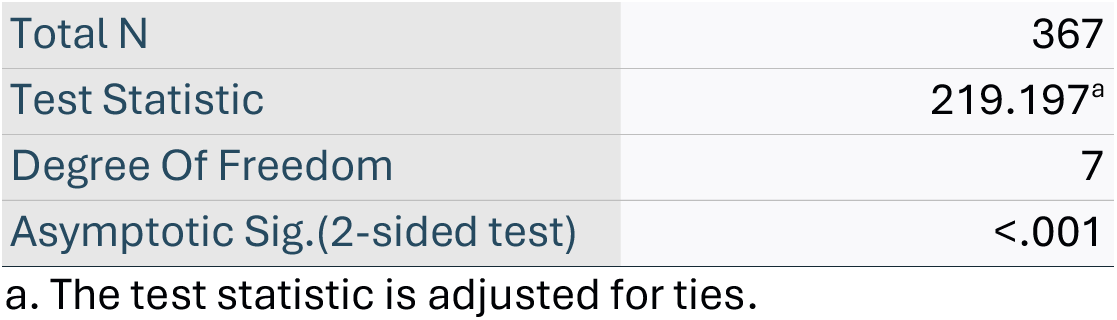

Pairwise Comparisons of Sites

| Sample 1-Sample 2 | Test Statistic | Std. Error | Std. Test<br>Statistic | Sig. | Adj. Sig.<sup>a</sup> |
| --- | --- | --- | --- | --- | --- |
| Tektite-Deep Tektite | 36.506 | 20.417 | 1.788 | .074 | 1.000 |
| Tektite-Europa | 38.956 | 20.417 | 1.908 | .056 | 1.000 |
| Tektite-Yawzi | -60.990 | 24.582 | -2.481 | .013 | .367 |
| Tektite-Grootpan | 133.363 | 20.495 | 6.507 | <.001 | .000 |
| Tektite-Booby Rock | 140.618 | 22.755 | 6.180 | <.001 | .000 |
| Tektite-Media Luna | 223.535 | 23.879 | 9.361 | <.001 | .000 |
| Tektite-San Cristobal | 249.801 | 23.095 | 10.816 | <.001 | .000 |
| Deep Tektite-Europa | -2.450 | 19.360 | -.127 | .899 | 1.000 |
| Deep Tektite-Yawzi | -24.483 | 23.711 | -1.033 | .302 | 1.000 |
| Deep Tektite-Grootpan | -96.856 | 19.442 | -4.982 | <.001 | .000 |
| Deep Tektite-Booby Rock | 104.112 | 21.811 | 4.773 | <.001 | .000 |
| Deep Tektite-Media Luna | -187.029 | 22.981 | -8.138 | <.001 | .000 |
| Deep Tektite-San Cristobal | -213.294 | 22.165 | -9.623 | <.001 | .000 |
| Europa-Yawzi | -22.033 | 23.711 | -.929 | .353 | 1.000 |
| Europa-Grootpan | -94.406 | 19.442 | -4.856 | <.001 | .000 |
| Europa-Booby Rock | 101.662 | 21.811 | 4.661 | <.001 | .000 |
| Europa-Media Luna | -184.579 | 22.981 | -8.032 | <.001 | .000 |
| Europa-San Cristobal | -210.844 | 22.165 | -9.512 | <.001 | .000 |
| Yawzi-Grootpan | 72.373 | 23.778 | 3.044 | .002 | .065 |
| Yawzi-Booby Rock | 79.628 | 25.751 | 3.092 | .002 | .056 |
| Yawzi-Media Luna | 162.545 | 26.749 | 6.077 | <.001 | .000 |
| Yawzi-San Cristobal | 188.811 | 26.052 | 7.248 | <.001 | .000 |
| Grootpan-Booby Rock | 7.255 | 21.883 | .332 | .740 | 1.000 |
| Grootpan-Media Luna | -90.173 | 23.050 | -3.912 | <.001 | .003 |
| Grootpan-San Cristobal | -116.438 | 22.237 | -5.236 | <.001 | .000 |
| Booby Rock-Media Luna | -82.917 | 25.081 | -3.306 | <.001 | .026 |
| Booby Rock-San Cristobal | -109.183 | 24.335 | -4.487 | <.001 | .000 |
| Media Luna-San Cristobal | -26.265 | 25.389 | -1.035 | .301 | 1.000 |

Each row tests the null hypothesis that the Sample 1 and Sample 2 distributions are the same. Asymptotic significances (2-sided tests) are displayed. The significance level is .050.
a. Significance values have been adjusted by the Bonferroni correction for multiple tests.

## Frontal Area

Generalized Linear model using Gamma distribution with a log link

Tests of Model Effects Type III
| Source | Wald Chi-Square | df | Sig. |
| --- | --- | --- | --- |
| (Intercept) | 981.683 | 1 | 0.000 |
| Site | 637.961 | 7 | 0.000 |

Dependent Variable: frontal\_area
Model: (Intercept), Site

Estimated Marginal Means

| Site | Mean | Std. Error | 95% Wald Confidence Interval |  |
| --- | --- | --- | --- | --- |
|  |  |  | Lower | Upper |
| Booby Rock | 0.2109376 | 0.02814119 | 0.1624034 | 0.2739762 |
| Deep Tektite | 0.1054058 | 0.01938339 | 0.0735080 | 0.1511452 |
| Europa | 0.0431872 | 0.00478910 | 0.0347508 | 0.0536719 |
| Grootpan | 0.1861410 | 0.02010056 | 0.1506345 | 0.2300168 |
| Media Luna | 1.2468598 | 0.17850110 | 0.9418011 | 1.6507301 |
| SanCristobol | 1.6289283 | 0.24060318 | 1.2194782 | 2.1758548 |
| Tektite | 0.0758304 | 0.01394469 | 0.0528827 | 0.1087361 |
| Yawzi | 0.0714491 | 0.01110447 | 0.0526871 | 0.0968921 |

Independent-Samples Kruskal-Wallis Test Summary

|  |  |
| --- | --- |
| Total N | 283 |
| Test Statistic | 198.124<sup>a</sup> |
| Degree Of Freedom | 7 |
| Asymptotic Sig.(2-sided test) | <.001 |

a. The test statistic is adjusted for ties.

Pairwise Comparisons of Sites

Pairwise Comparisons of Site
| Sample 1-Sample 2 | Test Statistic | Std. Error | Std. Test Statistic | Sig. | Adj. Sig.<sup>a</sup> |
| --- | --- | --- | --- | --- | --- |
| Europa-Tektite | -9.177 | 21.370 | -.429 | .668 | 1.000 |
| Europa-Yawzi | -10.684 | 18.999 | -.562 | .574 | 1.000 |
| Europa-Deep Tektite | 49.727 | 21.370 | 2.327 | .020 | .559 |
| Europa-Grootpan | -82.620 | 15.403 | -5.364 | <.001 | .000 |
| Europa-Booby Rock | 90.511 | 17.264 | 5.243 | <.001 | .000 |
| Europa-Media Luna | -177.721 | 18.020 | -9.862 | <.001 | .000 |
| Europa-SanCristobol | -194.424 | 18.380 | -10.578 | <.001 | .000 |
| Tektite-Yawzi | -1.507 | 23.960 | -.063 | .950 | 1.000 |
| Tektite-Deep Tektite | 40.550 | 25.880 | 1.567 | .117 | 1.000 |
| Tektite-Grootpan | 73.443 | 21.222 | 3.461 | <.001 | .015 |
| Tektite-Booby Rock | 81.334 | 22.608 | 3.598 | <.001 | .009 |
| Tektite-Media Luna | 168.544 | 23.191 | 7.268 | <.001 | .000 |
| Tektite-SanCristobol | 185.247 | 23.472 | 7.892 | <.001 | .000 |
| Yawzi-Deep Tektite | 39.043 | 23.960 | 1.629 | .103 | 1.000 |
| Yawzi-Grootpan | 71.936 | 18.833 | 3.820 | <.001 | .004 |
| Yawzi-Booby Rock | 79.827 | 20.383 | 3.916 | <.001 | .003 |
| Yawzi-Media Luna | 167.037 | 21.028 | 7.944 | <.001 | .000 |
| Yawzi-SanCristobol | 183.740 | 21.337 | 8.611 | <.001 | .000 |
| Deep Tektite-Grootpan | -32.893 | 21.222 | -1.550 | .121 | 1.000 |
| Deep Tektite-Booby Rock | 40.784 | 22.608 | 1.804 | .071 | 1.000 |
| Deep Tektite-Media Luna | -127.994 | 23.191 | -5.519 | <.001 | .000 |
| Deep Tektite-SanCristobol | -144.697 | 23.472 | -6.165 | <.001 | .000 |
| Grootpan-Booby Rock | 7.891 | 17.080 | .462 | .644 | 1.000 |
| Grootpan-Media Luna | -95.101 | 17.845 | -5.329 | <.001 | .000 |
| Grootpan-SanCristobol | -111.804 | 18.208 | -6.140 | <.001 | .000 |
| Booby Rock-Media Luna | -87.210 | 19.473 | -4.478 | <.001 | .000 |
| Booby Rock-SanCristobol | -103.913 | 19.807 | -5.246 | <.001 | .000 |
| Media Luna-SanCristobol | -16.703 | 20.470 | -.816 | .415 | 1.000 |

Each row tests the null hypothesis that the Sample 1 and Sample 2 distributions are the same. Asymptotic significances (2-sided tests) are displayed. The significance level is .050.
a. Significance values have been adjusted by the Bonferroni correction for multiple tests.

**Online Fig 2.**
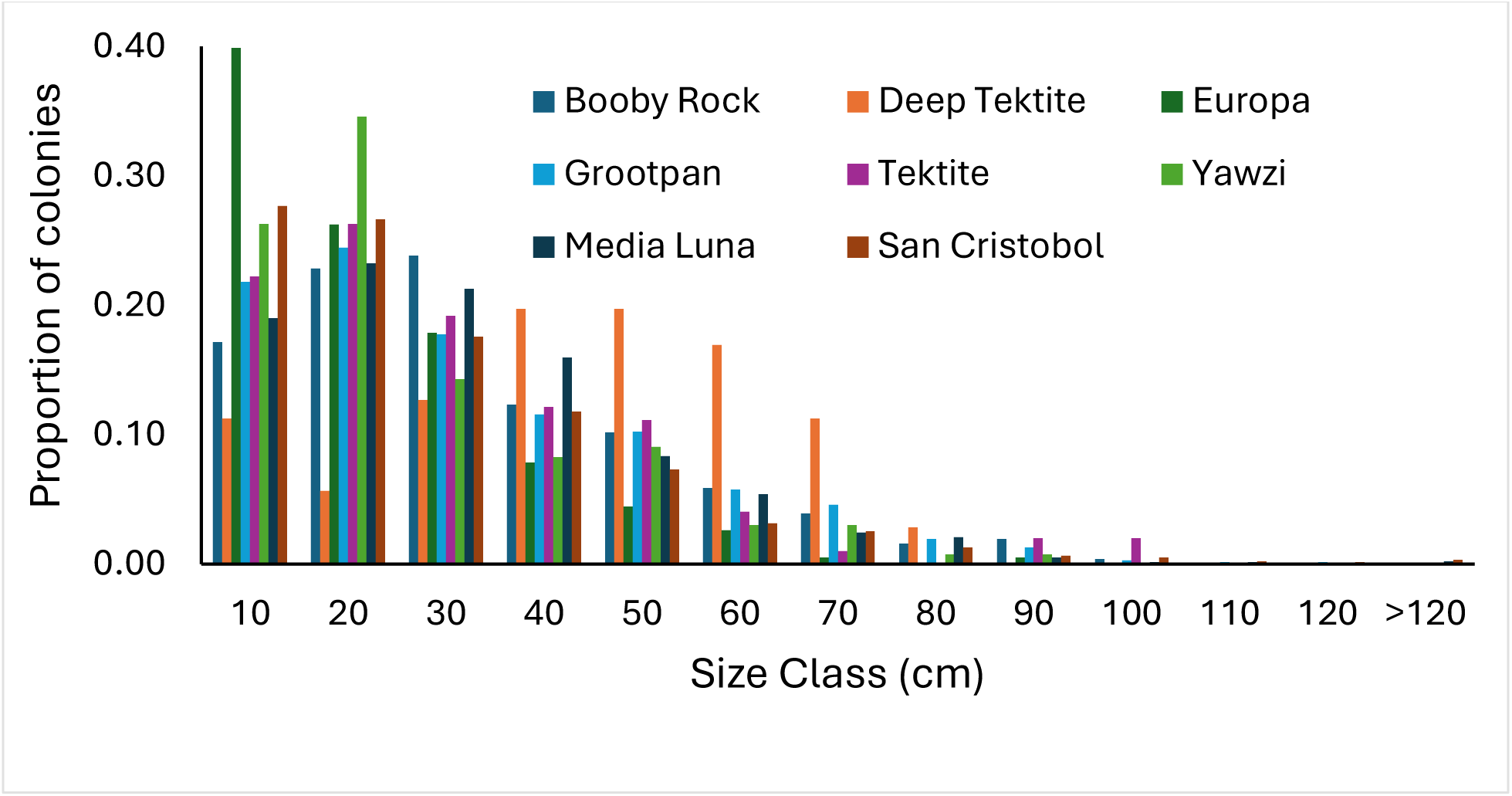
Size frequency distribution of octocoral colonies at 8 sites on Puerto Rico and St John.

### Comparison of canopy cover calculated from geometric models of colonies to and point counts from overhead images

The area of the canopy created by each colony was calculated as an ellipse using the derived breadth and thickness as the major and minor axes and those then summed to generate canopy area for each quadrat. A subset of the quadrats were imaged from overhead and those images were analyzed using CoralNet in which the perimeter of the colonies’ canopies was defined as a convex hull and the area within the canopy quantified from the proportion of 500 randomly placed points that were within the convex hull. Images from Puerto Rico were generated in 2022 at the same time as the census. St John images were also generated in 2022, which was 1 year after the census data were collected.

In general, there was good correlation between the two measures and either approach can distinguish high canopy cover quadrats from low canopy cover (Online Figure S3). However, differences between the estimates vary with the magnitude of cover. When the image based estimates were ranked, the geometric model approach overestimated canopy cover for 96% of the quadrats above the median value and underestimated cover for 65% of the quadrats below the median. Several of the estimates of canopy cover using the model approach were at or near 100% cover, which could occur if the canopy of different colonies overlapped. Canopy cover estimated from the images were not subject to that bias and the greatest cover estimate from the images was 52%. Additionally, the variance was considerably greater among the high octocoral density quadrats, primarily those from Puerto Rico. There was a significant relationship between the measures when the lower canopy cover sites were analyzed separately but no significant relationship among the high density sites. At high density there is likely overlap of canopies, reducing the counts and when in close proximity to each other, octocorals exhibit modified growth patterns making the characterization of their canopies as ellipses less accurate. Thus the canopy cover at high density will vary not only with the number and size of colonies, but also with their placement within the 1 m^2^ quadrat.

**Online Fig 3.**
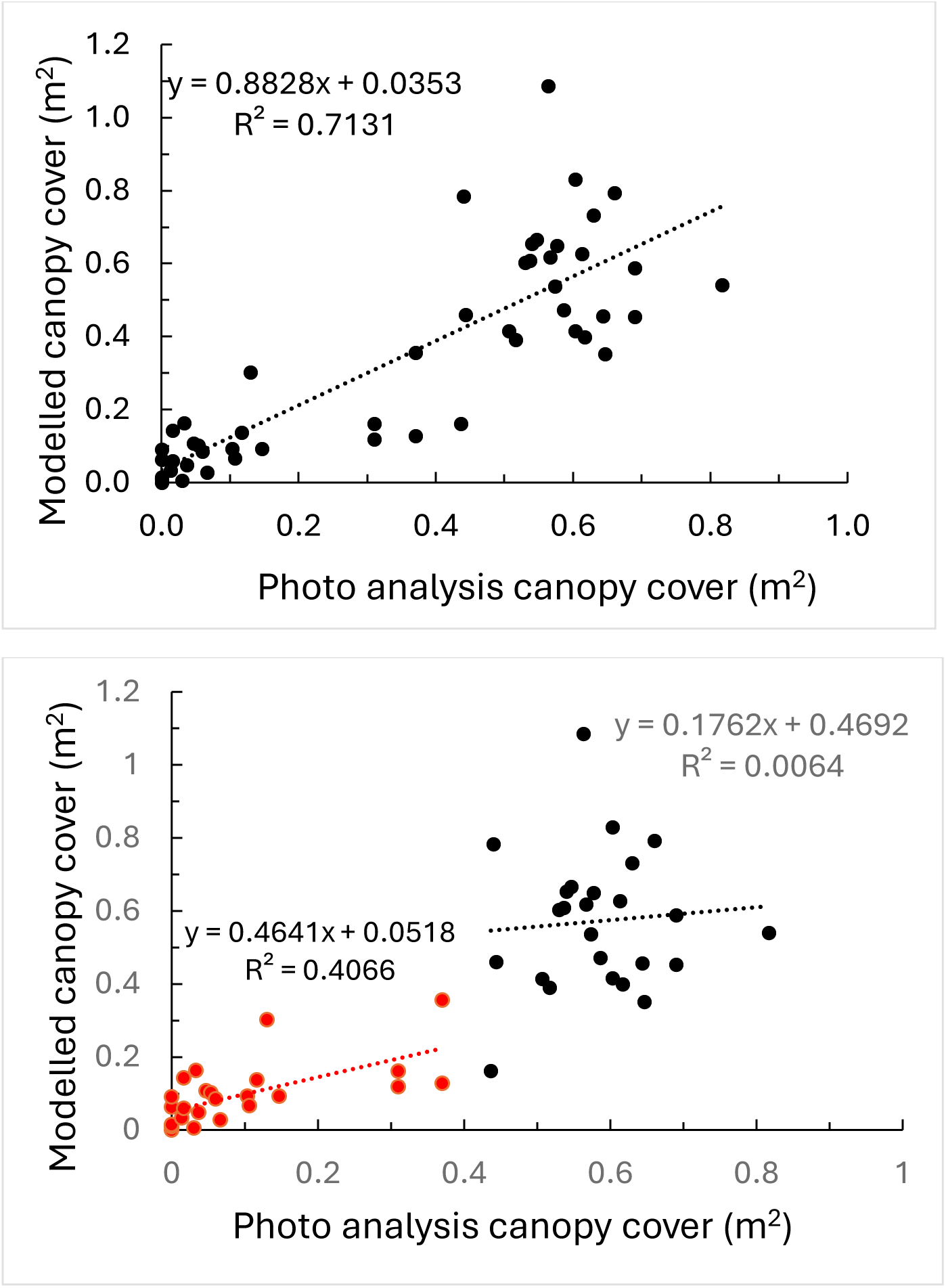
Comparison of canopy cover area calculated from colony heights (calculated canopy cover) and calculated from point counts of images of the quadrat. Upper, all quadrats. Lower, quadrats divided into those above (blue) and below (red) median canopy cover.

**Online Fig 4.**
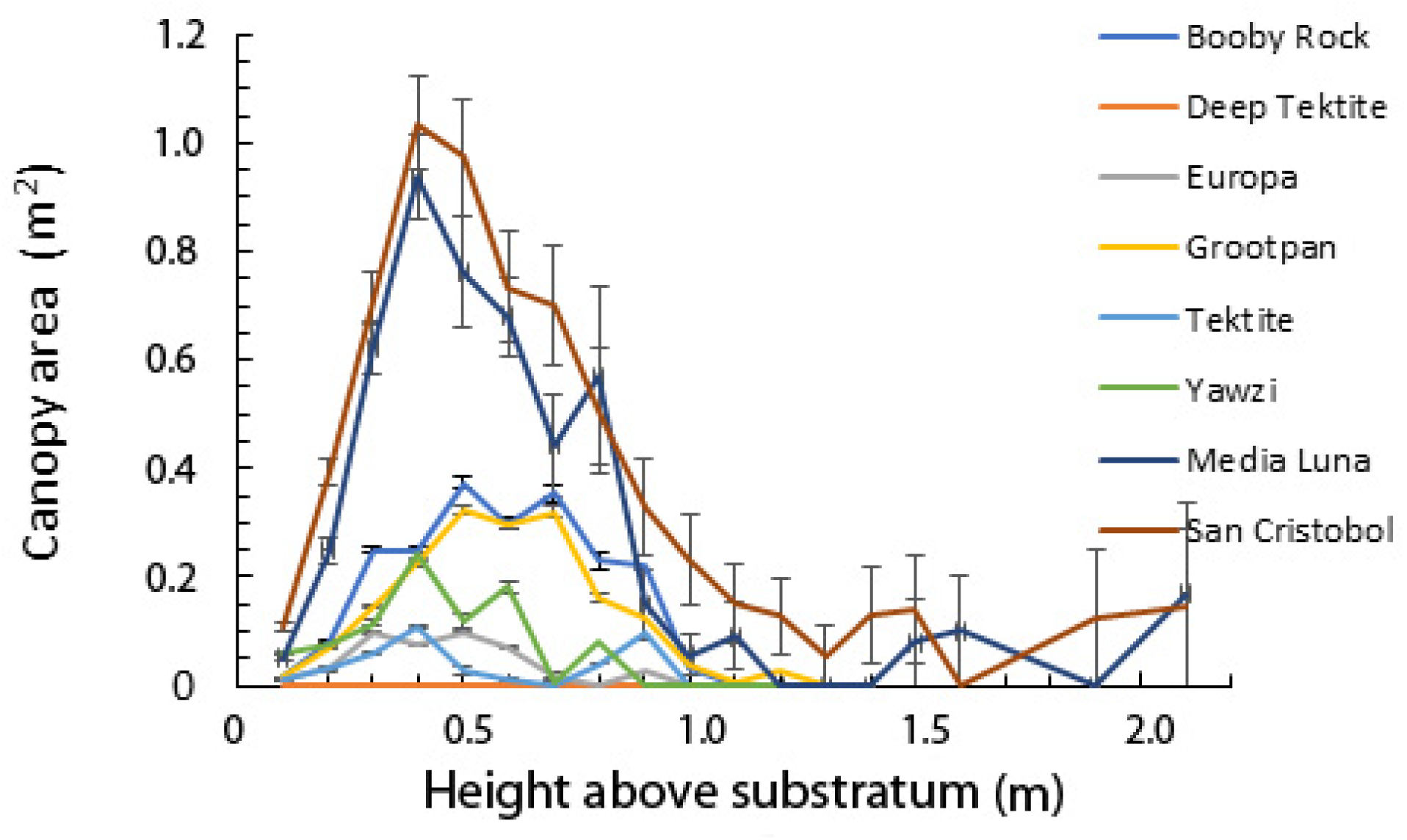
Canopy area relative to height above substratum. Values represent averages (+/- standard error) for quadrats at each study site.

**Online Fig 5.**
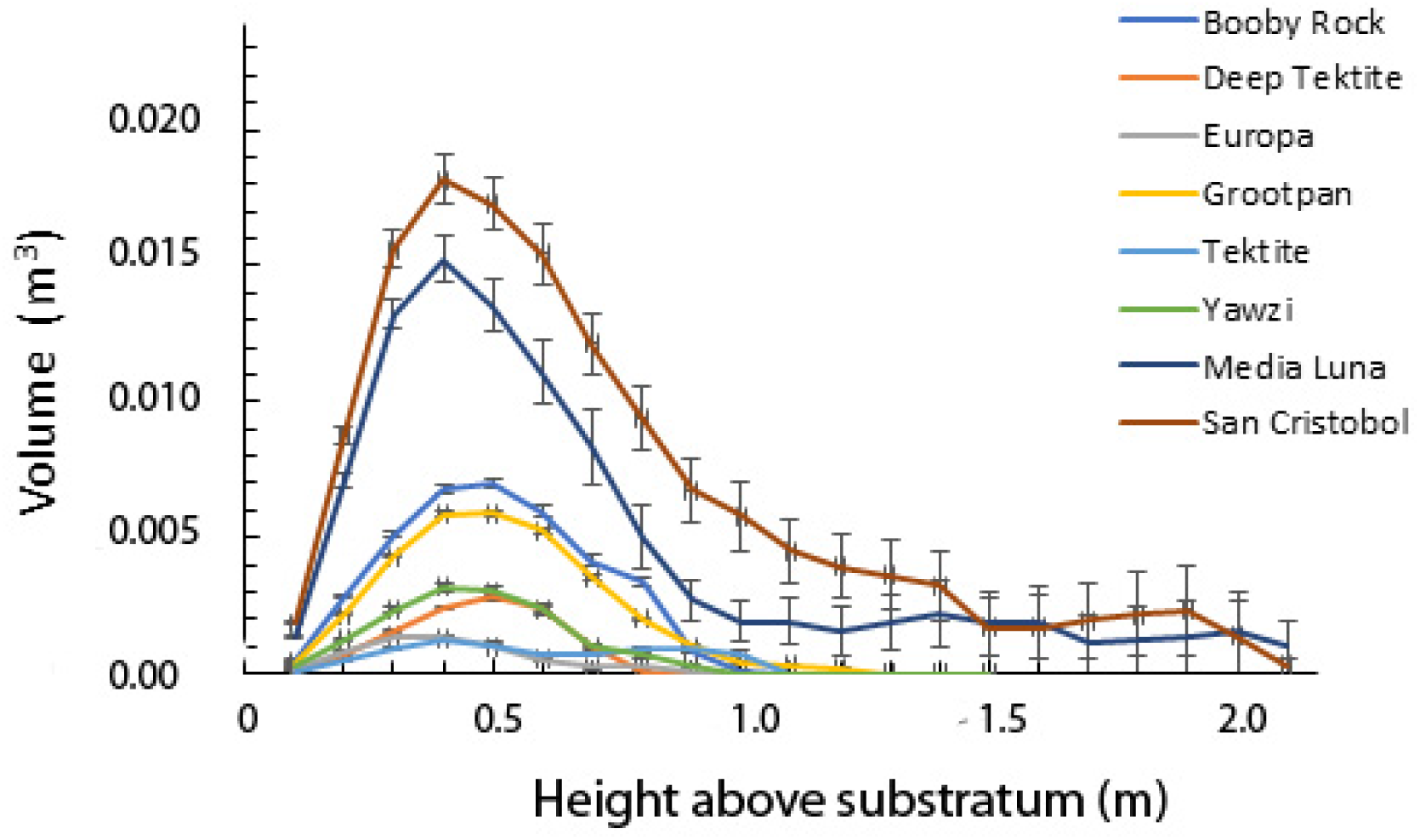
Average of summed volume (+/- standard error) occupied by colonies within 1 m^2^ areas, partitioned in 10 cm bands above the substratum. Values on the x-axis represent the upper edge of each band.

**Online Fig 6.**
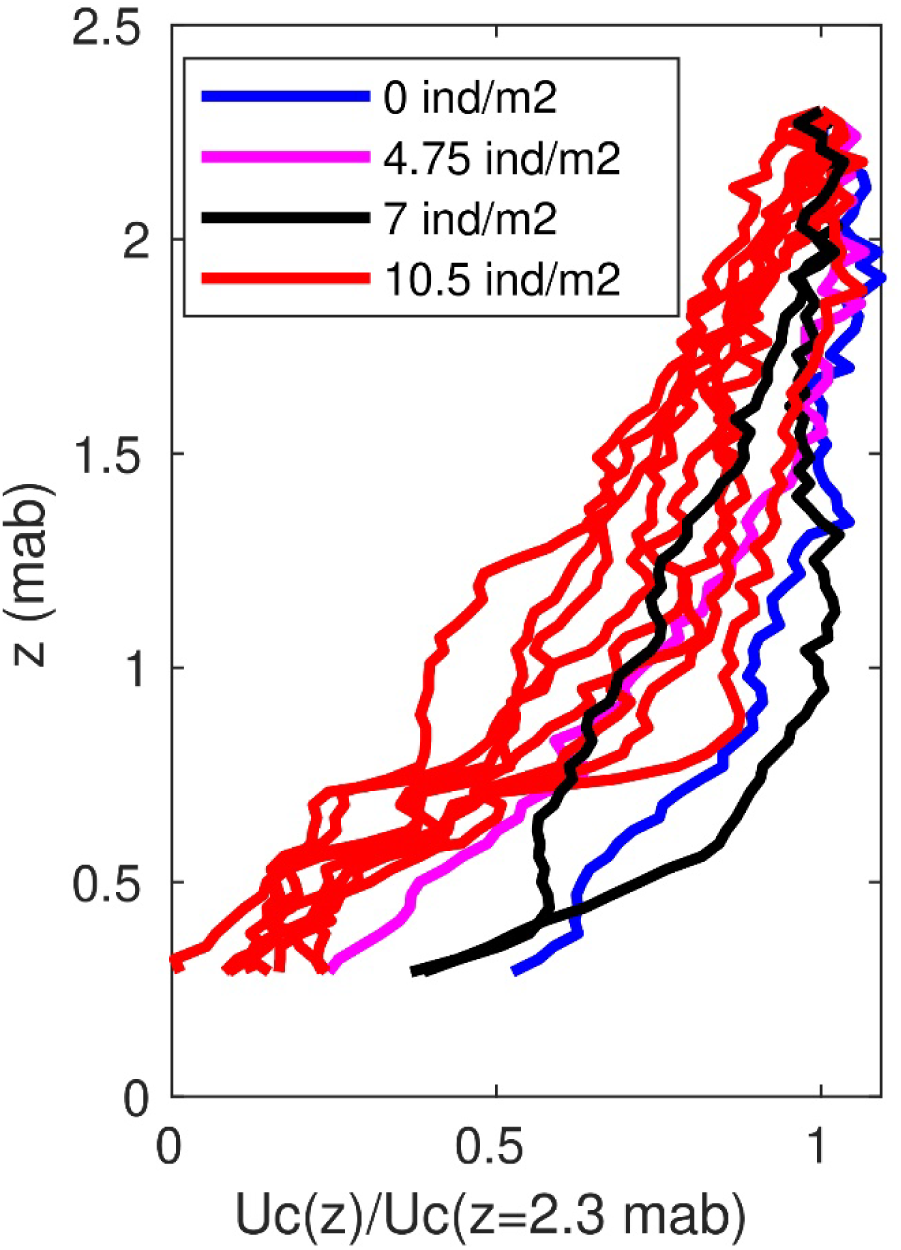
Vertical profile of mean flow speed (mean of 5 min flow velocity record) over a bare seabed and in three locations with octocorals population densities ranging from 4.75 to 10.5 individuals.m^-2^ on a fringing reef in Grootpan Bay, St John, US Virgin Islands. Mean flow speed is normalized to its value at 2.3 m above the bed.

## References

Abdolahpour M, Ghisalberti M, Lavery P, McMahon K (2017) Vertical mixing in coastal canopies. Limnology and Oceanography 62:26–42

Benayahu Y, Bridge TCL, Colin PL, Liberman R, McFadden CS, Pizarro O, Schleyer MH, Shoham E, Reijnen BT, Weis M, Tanaka J (2019) Octocorals of the Indo-Pacific

Bosch NE, Espino F, Tuya F, Haroun R, Bramanti L, Otero-Ferrer F (2023) Black coral forests enhance taxonomic and functional distinctiveness of mesophotic fishes in an oceanic island: implications for biodiversity conservation. Scientific Reports 13

Cerpovicz AF, Lasker HR (2021) Canopy effects of octocoral communities on sedimentation: modern baffles on the shallow-water reefs of St. John, USVI. Coral Reefs 40:295–303

Edmunds PJ, Tsounis G, Lasker HR (2016) Differential distribution of octocorals and scleractinians around St. John and St. Thomas, US Virgin Islands. Hydrobiologia 767:347–360

Falter JL, Atkinson MJ, Lowe RJ, Monismith SG, Koseff JR (2007) Effects of nonlocal turbulence on the mass transfer of dissolved species to reef corals. Limnology and Oceanography 52:274–285

FAO FaAOotUN- (2018) Global Forest Resources Assessment 2020-Terms and definitions Forest Resources Assessment Working Paper 188. United Nations, Rome

Gambrel B, Lasker HR (2016) Interactions in the canopy among Caribbean reef octocorals. Marine Ecology Progress Series 546:85–95

Gardner TA, Côté IM, Gill JA, Grant A, Watkinson AR (2003) Long-term region-wide declines in Caribbean corals. Science 301:958–960

Ghisalberti M, Nepf H (2006) The structure of the shear layer in flows over rigid and flexible canopies. Environ Fluid Mech 6:277–301

Girard JF, Edmunds PJ (2023) Effects of arborescent octocoral assemblages on the understory benthic communities of shallow Caribbean reefs. Journal of Experimental Marine Biology and Ecology 561:11

Jackson J, Donovan M, Cramer K, Lam V (2014) Status and trends of Caribbean coral reefs: 1970-2012. In: Jackson J, Donovan M, Cramer K, Lam V (eds), Washington, D.C. 306

Kupfner Johnson S, Hallock P (2020) A review of symbiotic gorgonian research in the western Atlantic and Caribbean with recommendations for future work. Coral Reefs 39:239–258

Lalas JAA, Jamodiong EA, Reimer JD (2024) Spatial patterns of soft coral (Octocorallia) assemblages in the shallow coral reefs of Okinawa Island, Ryukyu Archipelago, Japan: Dominance on highly disturbed reefs. Regional Studies in Marine Science 71

Lasker HR, Bramanti L, Tsounis G, Edmunds PJ (2020a) The rise of octocoral forests on Caribbean reefs. In: Riegl BM (ed) Advances in Marine Biology. Academic Press, pp361–410

Lasker HR, Martínez-Quintana A, Bramanti L, Edmunds PJ (2020b) Resilience of Octocoral Forests to Catastrophic Storms. Scientific Reports 10

Lenz EA, Bramanti L, Lasker HR, Edmunds PJ (2015) Long-term variation of octocoral populations in St. John, US Virgin Islands. Coral Reefs 34:1099–1109

Lowe RJ, Koseff JR, Monismith SG (2005) Oscillatory flow through submerged canopies: 1. Velocity structure. Journal of Geophysical Research-Oceans 110

Macdonald RW (2000) Modelling the mean velocity profile in the urban canopy layer. Bound-Layer Meteor 97:25–45

Monismith SG (2007) Hydrodynamics of coral reefs. Annual Review of Fluid Mechanics 39:37–55

Nelson H, Bramanti L (2020) From Trees to Octocorals: The Role of Self-Thinning and Shading in Underwater Animal Forests. In: Rossi S, Bramanti L (eds) Perspectives on the Marine Animal Forests of the World. Springer, Cham,

Nepf HM (2012) Flow and Transport in Regions with Aquatic Vegetation. In: Davis SH, Moin P (eds) Annual Review of Fluid Mechanics, Vol 44, pp123–142

Norstrom AV, Nystrom M, Lokrantz J, Folke C (2009) Alternative states on coral reefs: beyond coral-macroalgal phase shifts. Marine Ecology Progress Series 376:295–306

Orejas C, Carreiro-Silva M, Mohn C, Reimer JD, Samaai T, Allcock AL, Rossi S (2022) Marine Animal Forests of the World: Definition and Characteristics. Research Ideas and Outcomes 8:e96274

Otis N, Reimer JD, Kawamura I, Kise H, Mizuyama M, Obuchi M, Sommer B, McFadden CS, Beger M (2024) Variation in species and functional composition of octocorals and zoantharians across a tropical to temperate environmental gradient in the Indo-Pacific. Coral Reefs 43:613–626

Privitera-Johnson K, Lenz EA, Edmunds PJ (2015) Density-associated recruitment in octocoral communities in St. John, US Virgin Islands. Journal of Experimental Marine Biology and Ecology 473:103–109

Raupach MR, Finnigan JJ, Brunet Y (1996) Coherent eddies and turbulence in vegetation canopies: The mixing-layer analogy. Bound-Layer Meteor 78:351–382

Reichelt RE, Loya Y, Bradbury RH (1986) PATTERNS IN THE USE OF SPACE BY BENTHIC COMMUNITIES ON 2 CORAL REEFS OF THE GREAT-BARRIER-REEF. Coral Reefs 5:73–79

Rossi S, Bramanti L, Gori A, Orejas Saco del Valle C (2017) An overview of the animal forests of the world. In: Rossi S, Bramanti L, Gori A, Orejas Saco del Valle C (eds) Marine animal forests: The ecology of benthic biodiversity hotspots. Springer, pp1–26

Rossi S, Schubert N, Brown D, Soares MD, Grosso V, Rangel-Huerta E, Maldonado E (2018) Linking host morphology and symbiont performance in octocorals. Scientific Reports 8

Ruzicka RR, Colella MA, Porter JW, Morrison JM, Kidney JA, Brinkhuis V, Lunz KS, Macaulay KA, Bartlett LA, Meyers MK, Colee J (2013) Temporal changes in benthic assemblages on Florida Keys reefs 11 years after the 1997/1998 El Nino. Marine Ecology Progress Series 489:125–141

Sanchez JA, Gomez-Corrales M, Gutierrez-Cala L, Vergara DC, Roa P, Gonzalez-Zapata FL, Gnecco M, Puerto N, Neira L, Sarmiento A (2019) Steady Decline of Corals and Other Benthic Organisms in the SeaFlower Biosphere Reserve (Southwestern Caribbean). Frontiers in Marine Science 6

Tsounis G, Edmunds PJ (2017) Three decades of coral reef community dynamics in St. John, USVI: a contrast of scleractinians and octocorals. Ecosphere 8

Tsounis G, Edmunds PJ, Bramanti L, Gambrel B, Lasker HR (2018) Variability of size structure and species composition in Caribbean octocoral communities under contrasting environmental conditions. Marine Biology 165

Umanzor S, Ladah L, Zertuche-González JA (2018) Intertidal Seaweeds Modulate a Contrasting Response in Understory Seaweed and Microphytobenthic Early Recruitment. Frontiers in Marine Science 5

Velásquez J, Sánchez JA (2015) Octocoral Species Assembly and Coexistence in Caribbean Coral Reefs. Plos One 10

Wells CD, Muñoz-Maravilla JD, Lasker HR, Edmunds PJ (2022) Information legacies of early ecological studies. Bulletin of Marine Science 98:493–494

Williams SM, Mumby PJ, Chollett I, Cortes J (2015) Importance of differentiating Orbicella reefs from gorgonian plains for ecological assessments of Caribbean reefs. Marine Ecology Progress Series 530:93–101

Yoshioka PM, Yoshioka BB (1989) Effects of wave energy, topographic relief and sediment transport on the distribution of shallow-water gorgonians of Puerto Rico. Coral Reefs 8:145–152

Yranzo A, Villamizar E, Romero M, Boadas H (2014) Structure of the coral and octocoral communities of Isla de Aves, Venezuela, Northeast Caribbean. Revista De Biologia Tropical 62:115–136

